# The promise and challenge of spatial inference with the full ancestral recombination graph under Brownian motion

**DOI:** 10.1101/2024.04.10.588900

**Authors:** Puneeth Deraje, James Kitchens, Graham Coop, Matthew M. Osmond

**Affiliations:** Department of Ecology & Evolutionary Biology, University of Toronto; Department of Evolution & Ecology and Center for Population Biology, University of California - Davis

**Keywords:** Ancestral recombination graph, spatial population genetics, population genetic inference, Brownian motion, networks, genetic ancestry

## Abstract

Spatial patterns of genetic relatedness among samples reflect the past movements of their ancestors. Our ability to untangle this history has the potential to improve dramatically given that we can now infer the ultimate description of genetic relatedness, the ancestral recombination graph (ARG). By extending spatial theory previously applied to trees, we generalize the common model of Brownian motion to full ARGs, thereby accounting for correlations in trees along a chromosome while efficiently computing likelihood-based estimates of dispersal rate and genetic ancestor locations, with associated uncertainties. We evaluate this model’s ability to reconstruct spatial histories using individual-based simulations and unfortunately find a clear bias in the estimates of dispersal rate and ancestor locations. We investigate the causes of this bias, pinpointing a discrepancy between the model and the true spatial process at recombination events. This highlights a key hurdle in extending the ubiquitous and analytically-tractable model of Brownian motion from trees to ARGs, which otherwise has the potential to provide an efficient method for spatial inference, with uncertainties, using all the information available in the full ARG.

## 1 Introduction

Life moves – offspring disperse, populations admix, and species’ ranges shift. While most of these events go unnoticed in the moment, they leave spatial patterns in the genetic diversity of a sample. Though often faint, we can use such signals to gain insight into the spatial history of the samples’ shared genetic ancestors.

One broad set of approaches to infer spatial history divides sample genomes up into a small number of geographic regions and, using allele frequency (e.g., Excoffier *et al*., 2021) or gene trees (e.g., Müller *et al*., 2018), estimates the split times and rates of gene flow between these regions. While useful, these approaches require the *a priori* grouping of samples and can obscure the fact that there can be population structure at many geographical scales. For example, in many species genetic differentiation builds up relatively smoothly with the geographic distance between samples. Such patterns motivate modeling approaches that treat space explicitly in so-called isolation-by-distance models. A subset of these models assume local migration between groups of individuals (demes) on a lattice (e.g., the stepping-stone model, Kimura and Weiss, 1964; Malécot, 1948), allowing the rate of increase in genetic differentiation (*F*_*ST*_) with geographic distance to be used to estimate dispersal rates (Rousset, 2000, 1997). The alternative is to treat space in a truly continuous manner, avoiding the need to assume well-mixed demes or a fixed layout of individuals. At its core, the classic continuum model (Malécot, 1948; Wright, 1943) assumes lineages move as independent Brownian motions. The tractability of this model makes it useful for spatial inference, for example, inferring dispersal rate and the locations of genetic ancestors from gene trees (e.g., Lemmon and Lemmon, 2008; Novembre and Slatkin, 2009). As a result, independent Brownian motion remains a common feature of many phylogeographic methods (e.g., Dellicour *et al*., 2021). Further research has aimed at resolving the issues that arise when assuming lineages move as independent Brownian motions, namely the clustering of individuals in forward-in-time models (Felsenstein, 1975) and sampling inconsistency in backward-in-time models (Barton *et al*., 2010a). The spatial Λ-Fleming-Viot process (Barton *et al*., 2010a, 2013, 2010b) is an alternative that, among other things, incorporates local density dependence to avoid these issues. However, the model’s mathematical complexity makes inference computationally expensive (Wirtz and Guindon, 2023) and therefore limited to small sample sizes. Thus, independent Brownian motion, despite its limitations, continues to be an analytically tractable and computationally feasible model that is often useful for spatial inference, at least when dealing with non-recombining sequences.

On the empirical front, the increasing feasibility of whole-genome sequencing has led to an influx of genetic data and motivated advances in the inference of the genealogical history of a sample undergoing recombination (e.g., Deng *et al*., 2024; Kelleher *et al*., 2019; Rasmussen *et al*., 2014; Schaefer *et al*., 2021; Speidel *et al*., 2019; Wohns *et al*., 2022). Recombination allows different regions of the same chromosome to have different gene trees. Although single-tree approaches may be suitable when studying non-recombining sequences (e.g., mitochondrial DNA), they do not capture the range of genetic relationships found across recombining genomes nor the correlations between these relationships. For this, we must turn to the ancestral recombination graph (ARG).

An ARG encodes the complete genetic history of a sample of recombining genomes (Griffiths and Marjoram, 1996; Hudson, 1983; Lewanski *et al*., 2023). As such, it has proven to be an incredibly rich source of information about the history of the sample (Harris, 2019; Hejase *et al*., 2020; Lewanski *et al*., 2023). One particularly promising application is spatial inference. Two recent methods use an ARG to provide point estimates of ancestor locations (Grundler *et al*., 2024; Wohns *et al*., 2022) while another provides the full probability distribution for an ancestor’s location but ignores correlations between trees (Osmond and Coop, 2024). Here, we extend the classic Brownian motion model for trees to describe movement down an ARG. Under this model, we derive the full likelihood of the sample locations given an ARG, allowing us to infer the dispersal rate and the probability distribution for the location of every genetic ancestor in the ARG using the complete genealogical history of the sample. In doing so, we highlight a key problem in extending a classic model of movement from trees to ARGs.

## 2 Brownian motion on an ARG

An ARG is a graphical representation of the genealogical history of a set of sample genomes that may have undergone recombination in the past (Wong *et al*., 2023). As the history of the samples at each site in the genome can be depicted as a tree, an ARG weaves together these histories based upon their shared structure. Altogether, an ARG provides the complete genetic history of the samples across the genome.

Each node in an ARG represents a haploid genome. Edges are directed from an ancestral node to a descendant node, describing the line of inheritance over time. A node that is the product of recombination has two ancestral nodes with annotated edges referring to the specific regions of the genome that were inherited from each. Time is measured backwards from the present, starting from the most recent sample node at time *t*= 0 and increasing as we go deeper into the past towards the root of the ARG, its grand most recent common ancestor (GMRCA). In this work we focus on “full ARGs” (Wong *et al*., 2023), which include the complete set of coalescent, recombination, and common ancestor nodes. This is mainly for ease of explanation. Our conclusions and algorithms are applicable more broadly to commonly inferred “simplified ARGs”, which only include coalescent nodes found in the local trees (Baumdicker *et al*., 2022; Lewanski *et al*., 2023; Shipilina *et al*., 2023; Wong e*t al*., 2023).

We model the movement of genetic material forward in time by Brownian motion with dispersal rate *σ*^2^ (see Table 1 for a list of key symbols). In other words, we assume that the location of a node is normally distributed about its parent node with variance *σ*^2^*t*, where *t*is the length of the edge connecting them (in generations). In each generation, autosomal inheritance is equally likely via the mother or father; therefore, the effective variance is the average of maternal and paternal variances (e.g., Smith *et al*., 2023). While the computations are shown for one dimension, they are readily extended to two dimensions by replacing the dispersal rate, *σ*^2^, with a dispersal matrix, 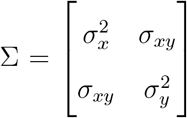, where *σ*_*xy*_ is the covariance in offspring locations across the two dimensions.

**Table 1:** Description of key symbols used.

| Symbol | Name | Description (and notes) |
| --- | --- | --- |
| $n_s$ | Number of samples | |
| $n_p$ | Number of paths | Total number of paths in the ARG from a sample to a root |
| $n_r$ | Number of roots | |
| $\sigma^2$ | Dispersal rate | Variance in offspring locations around parent |
| $\mathbf{S}_p$ | Path matrix | Shared times between each pair of paths (could refer to either the full path matrix or the minimal path matrix) |
| $\mathbf{S}$ | Sample matrix | $\sigma^2 \mathbf{S}$ is covariance matrix between sample locations |
| $\mathbf{P}$ | Path-sample matrix | $n_p \times n_s$ matrix whose entry is 1 if path $i$ is associated with sample $j$ |
| $\mathbf{R}$ | Path-root matrix | $n_p \times n_r$ matrix whose entry is 1 if path $i$ is associated with root $j$ |
| $\vec{\ell}^*$ | Observed sample locations | Vector of length $n_s$ |
| $\vec{L}$ | Sample location vector | Vector-valued random variable of sample locations |
| $\vec{L}_p$ | Path location vector | Vector-valued random variable of sample locations at the end of each path |
| $\vec{\mu}$ | Root location vector | Vector of length $n_r$ |
| $L_a$ | Internal node location | Random variable denoting location of internal node $a$ |
| $\vec{s}_a$ | | Vector-valued random variable with shared time between the path of internal node $a$ and each path in $\mathbf{S}_p$ |
| $V$ | Ancestor specific uncertainty | Component of location variance related to the position of the ancestor in the ARG and root uncertainty (not dispersal) |
| $t_a$ | | Time from node $a$ to the root |

We assume we know the ARG with certainty and therefore all edges are treated equally regardless of the amount of genome they span; if we know with certainty that an edge exists in the graph, we do not care how much genetic material those ancestors contributed to the samples. This differs from the approach of Grundler *et al*. (2024), which gives less weight to edges with smaller spans due to the lower confidence in accurately inferring these short span edges from genetic data. Here we are more focused on developing the underlying theory rather than pragmatically applying a method.

All of the equations presented below are generalizations of the standard treebased approach to account for recombination. As a tree is simply an ARG without recombination, these generalized equations will all collapse back to those previously presented in literature for a single tree.

The first step in estimating dispersal rates and the locations of genetic ancestors given an ARG is calculating a likelihood for the sample locations. Under Brownian motion, the vector of sample locations, specifically, 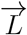, has a multivariate normal distribution, specifically,

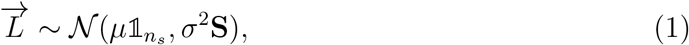

where *μ* is the location of the root (GMRCA) of the ARG, 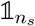 is a column vector of length *n*_*s*_ (number of samples) consisting of just ones, *σ*^2^ is effective the dispersal rate, and *σ*^2^**S** is the covariance matrix between the sample locations. We refer to **S** as the sample matrix and it represents the covariance between the samples that is solely due to the structure of the ARG, independent of the dispersal rate.

For a single tree we can assume that the displacement along any edge is independent of the other edges’ displacements. The displacement of an edge from node *i* to node *j* is distributed as *D*_*i,j*_ ∼ *𝒩* (0, *σ*^2^*t*_*i,j*_), where *t*_*i,j*_ is the length of the edge in generations. A sample’s location is given by the sum of the displacements along the edges from the root to the sample. We refer to a series of edges connecting an older (further in the past) node to a more recent node as a path. We are particularly interested in paths from the root to a sample and call the set of such paths the sample paths. The covariance between the locations of two samples is then simply *σ*^2^ times the shared time between the unique corresponding sample paths, hence the entries of **S** are just the shared time between each pair of samples, as mentioned above.

In generalizing this approach to an ARG we run into two related issues. First, each recombination node creates a loop within the graph. This means that samples found below a recombination node will have multiple paths back to the root. Therefore, computing the entries of **S** for an ARG is not as straightforward as it is for a tree and the covariance between samples must take into account this braided history where multiple paths lead to the same sample. Second, the presence of these loops means that we can no longer always assume that the displacements along edges of the graph forward in time are independent. Assuming that the parents of a recombination node must precisely meet each other in space, the displacement along the edges involved in the loop must satisfy the condition that the sum of the displacements around the left half of the loop equal the sum around the right half. For example, in Figure 1 the displacement from node *G* to node *F* plus the displacement from node *F* to node *E* must equal the displacement from node *G* directly to node *E, D*_*G,F*_ + *D*_*F,E*_ = *D*_*G,E*_. We refer to the collection of these conditions across an ARG as *η*_loops_.

**Figure 1:**
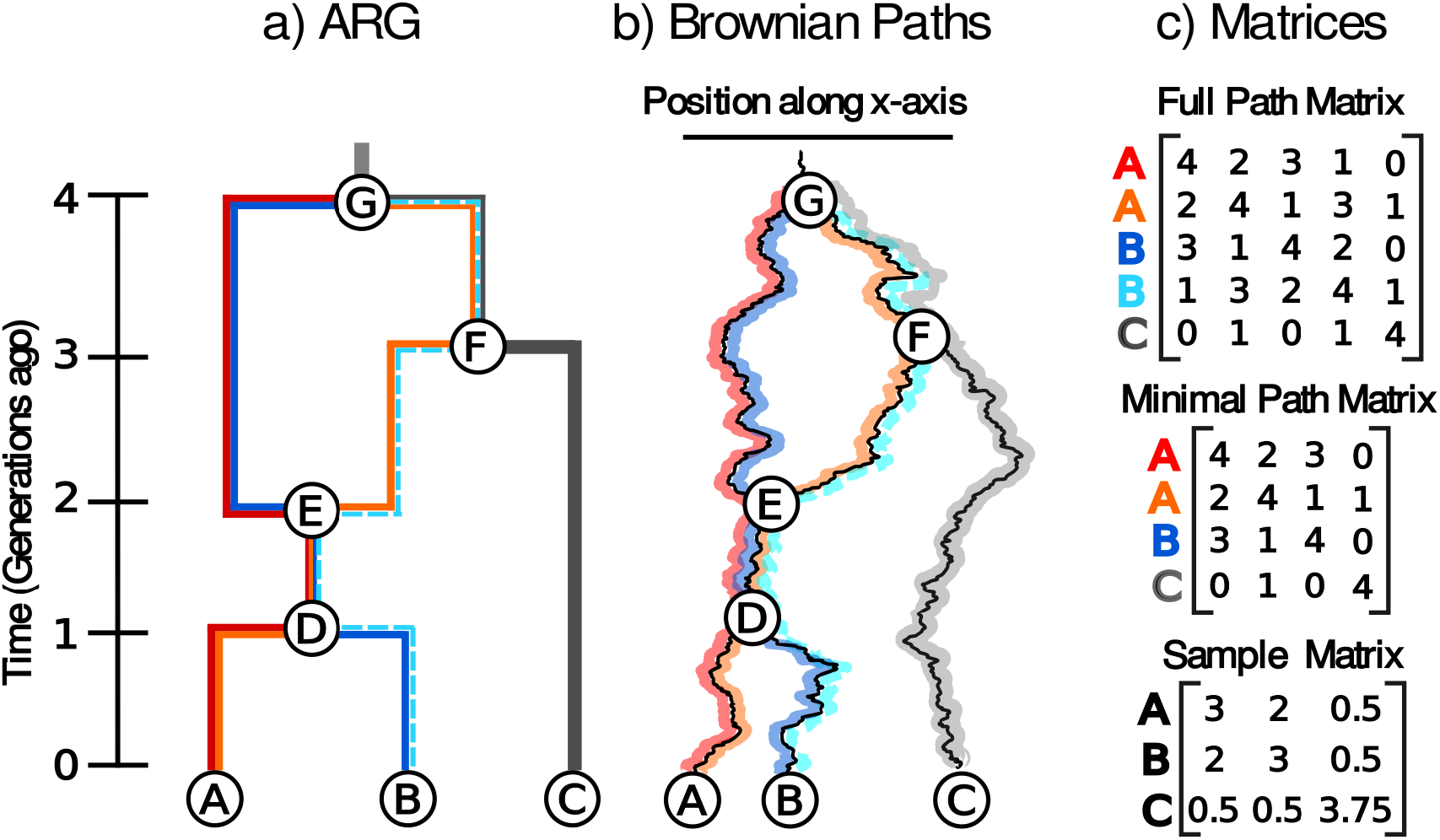
Brownian motion on an ARG. (a) An ARG, (b) an instance of Brownian motion along the ARG, and (c) the corresponding path (**S**_*p*_) and sample (**S**) matrices. Sample paths through the ARG are individually colored. Samples *A* and *B* are associated with two paths each - red/orange and navy/sky blue, respectively - because they are beneath a recombination node (node *E*). In the Brownian motion model illustration, note that the paths meet precisely at the recombination node *E*. The path matrices contain the shared time between the sample paths. The sky blue path from sample B is excluded when calculating the minimal path matrix as it provides redundant information.

To account for these conditions, *η*_loops_, we start by assuming (incorrectly) that Brownian motion along each edge of the ARG is indeed independent, as in the case of trees. This leads to a sample node having different location distributions based on the choice of sample path. Nevertheless, we encode the distribution of locations of the nodes at the tips of each sample path in 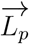, a column vector of length *n*_*p*_ (number of sample paths) with distribution

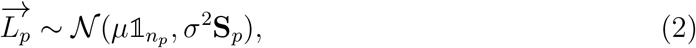

where **S**_*p*_, referred to as the path matrix, is the matrix of shared times between sample paths. To get the correct distribution of sample locations, 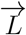 (Equation 1), we need to condition 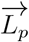 so that all sample paths that end at the same sample also end at the same location. We refer to this condition as *η*_paths_ and show that *η*_paths_ reduces to *η*_loops_ (Section S2). The sample matrix can then be easily computed from the path matrix (Section S1),

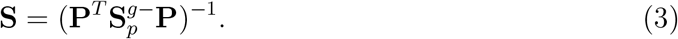

Here, 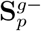 is the generalized inverse of **S**_*p*_ and **P** is a conversion matrix of size *n*_*p*_ *× n*_*s*_ that pairs each sample path *i* with its associated sample *j*. The *i, j*^*th*^ element of **P** is 1 if path *j* starts at sample *i*, otherwise it is 0. As a check, note that the sample matrix and path matrix are equivalent for a tree as each sample is associated with only one path.

We can now compute the likelihood of sample locations (Equation 1). Though this conversion from **S**_*p*_ to **S** is possible, it is generally unnecessary and it is anyways easier to work with the path matrix in what follows. Additionally, while we could calculate the shared times between every pair of sample paths through the ARG, many of the paths are not linearly independent from one another and are therefore redundant in our calculations. In practice, we use only a minimal subset of paths and its corresponding minimal path matrix (see Figure 1 for an example). While the total number of sample paths grows faster than linearly with the number of recombination events, the minimal number of sample paths grows linearly (being equal to the number of samples plus the number of recombination events; see Section S3). We have developed an algorithm to calculate the minimal path matrix in a single tip-to-root traversal of the ARG (Section S3).

### 2.1 Multiple roots

For multiple reasons we might want to focus on just the recent past. First, long-term movement may not be accurately captured by Brownian motion due to geographic boundaries or large-scale population movements (Ianni-Ravn *et al*., 2023). Second, as we move further back into the past, sample lineages can become spatially well mixed (Wakeley, 1999), which makes it difficult or impossible to extract meaningful spatial information about the deep history of samples. Third, with ARG inference, deeper nodes in the ARG are often poorly resolved, both in timing and topology (e.g., see Figure S2B in Fan *et al*., 2023). To avoid these issues, we will often want to cut off an ARG at a given time in the past and ignore any deeper connections.

When we chop an ARG below the GMRCA the graph no longer has a single root (Figure S1). Instead there are multiple roots, i.e., nodes with no parent nodes, each associated with specific paths through the ARG. Let 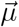 be the vector of root locations (of length *n*_*r*_, the number of roots) and **R** be a conversion matrix of size *n*_*p*_ *× n*_*r*_. The *i, j*^*th*^ element of **R** is 1 if path *j* terminates at root *i*, otherwise it is 0. Then 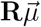 is the vector of root locations associated with each of the *n*_*p*_ paths.

The full covariance between the locations of the samples at the ends of the sample paths is the covariance created by the structure of the ARG plus the covariance between the root locations, 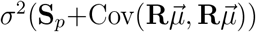. Here, we assume that the root locations are independent of each other and that each has zero variance, 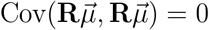, so that the covariance between the locations of the samples at the ends of the sample paths remains *σ*^2^**S**_*p*_. The assumption of independence is reasonable if we cut off the ARG at a point by which the ancestors are well mixed in a finite habitat. The assumption of zero variance in root locations can be relaxed by adding a variance to the covariance term between pairs of paths starting at the same root. The resulting likelihood of sample locations is

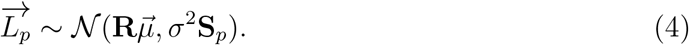

### 2.2 Estimates

If 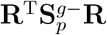 is invertible, we can compute the MLEs for the root locations given the observed sample locations, 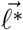, as the unique solution to (Section S1.2)

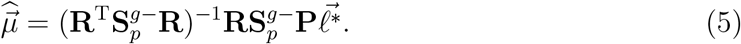

Once we have estimated the locations of the roots, we can compute the MLE of the dispersal rate,

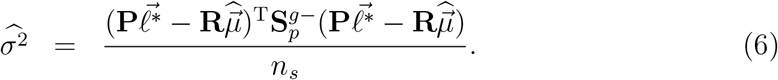

With estimates of both the root locations and the dispersal rate, we can then calculate the distribution of any internal node location (a genetic ancestor). Given an internal node, we choose an arbitrary path from that node to one of the roots. Conditional on the dispersal rate, root locations, and observed sample locations, the location of an internal node, *L*_*a*_, is normally distributed with mean

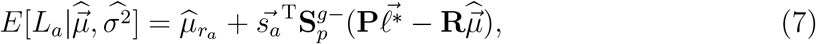

where 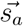 is the vector of shared times with the minimal paths used to construct **S**_*p*_ and 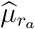 is the MLE location of the *r*_*a*_ root (the *r*_*a*_^th^ element of 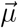). The total variance in the internal node’s location is a combination of the variance due to Brownian motion and the variance due to uncertainty in the root locations,

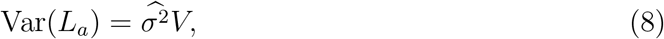

Where

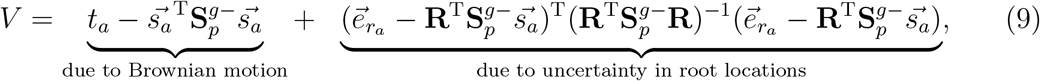

and 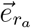 is a length *n*_*r*_ column vector that is zero everywhere except with a 1 at the *r*_*a*_^th^ position.

## 3 Building intuition

### 3.1 Ancestor locations

Changing either the topology or branch lengths within an ARG will in general change the dispersal rate and ancestral location estimates. We show the latter in Figure 2, where we compare against two previous approaches: 1) the MLE location with Brownian motion on each marginal tree (as in Osmond and Coop, 2024) and 2) simple averaging, where a node’s location is the average of its child nodes’ locations (as in Wohns *et al*., 2022, except we use the full ARG instead of the simplified one). By separately altering the timing of various nodes in the graph (*E, F*, and *G*), we observe a change in the MLE location of node *H* under our model. In contrast, point estimates from the averaging method are unaffected by changes to node timing (green lines in Figure 2b), as this approach does not incorporate edge lengths. Meanwhile, treebased MLEs (red and blue curves in Figure 2b) respond to changes in the timing of a node only if it affects the edge lengths in the given tree (e.g., the timing of node *G* only affects shared times in the blue tree). Recombination nodes (e.g., node *F*) do not affect the shared times in trees and so their timing does not affect tree-based estimates. When the timing of a node affects a tree and is in a loop (e.g., node *G*), the ARG and tree estimates can show opposite trends (black vs. blue curves in Figure 2b-iii). Note that under a model of Brownian motion, moving node *G* further into the past should move the location of node *H* towards the locations of nodes *A* and *B* and away from the location of node *C* (as it allows more time for the node *C* to wander away). This behavior is only captured when using Brownian motion on the ARG, not just on an individual tree.

**Figure 2:**
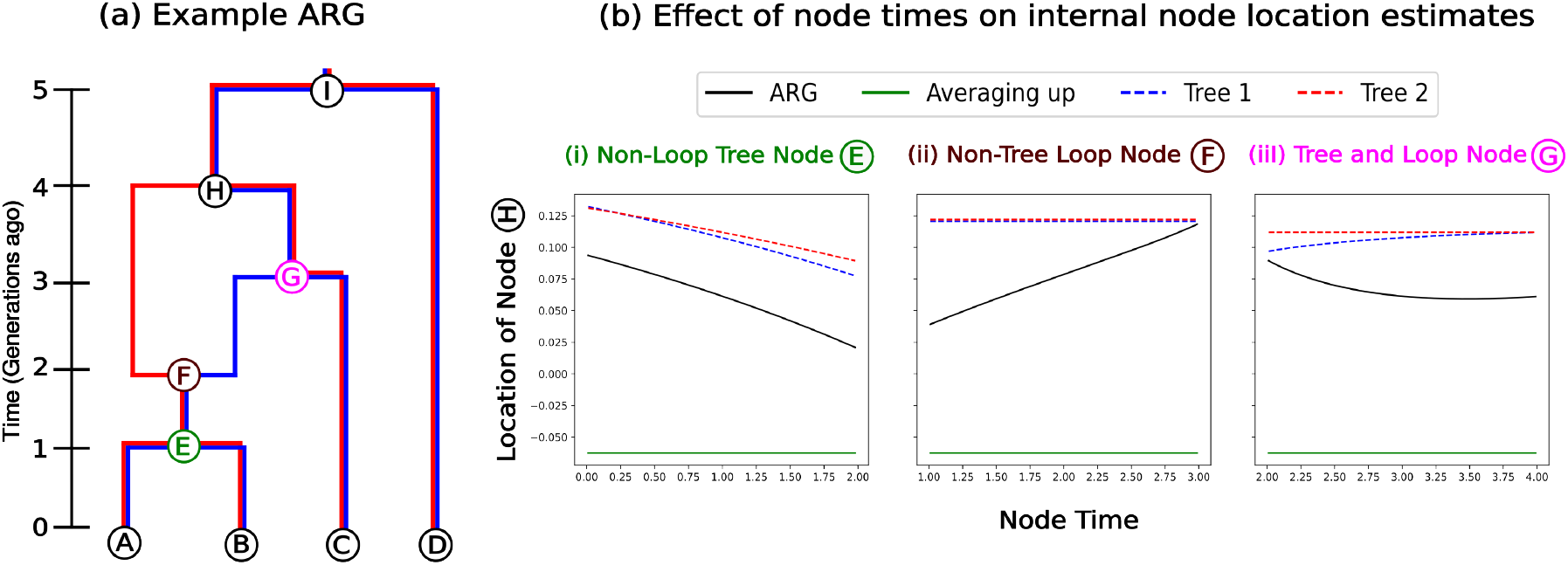
Ancestor locations. (a) Toy 2-tree ARG. (b) Most likely location of node *H* computed using the ARG (black), individual trees (red and blue), and the averaging-up method (green), plotted as a function of the time of (i) a node that alters individual trees and the ARG but is not part of the ARG loop, (ii) a node that doesn’t alter individual trees but is part of the ARG loop and (iii) a node which is part of the ARG loop that alters a tree and the ARG. Locations of nodes *A, B, C* and *D* are −0.5, 0, 0.5 and 1, respectively.

With the greater amount of information contained within the ARG vs. a single tree, we always estimate ancestor locations with higher certainty than the tree-based method provided we use the same dispersal rate (Figure 3c). In the absence of any other information, the variance in the location estimate increases linearly with time under Brownian motion. Both coalescent and recombination nodes bring additional information and hence reduce the increase in uncertainty of the location estimates as the lineage is tracked backwards in time (Figure 3c). Larger ARGs (either due to more samples or more trees) would have more such nodes bringing in extra information and so would show a greater reduction in variance compared to a single tree.

**Figure 3:**
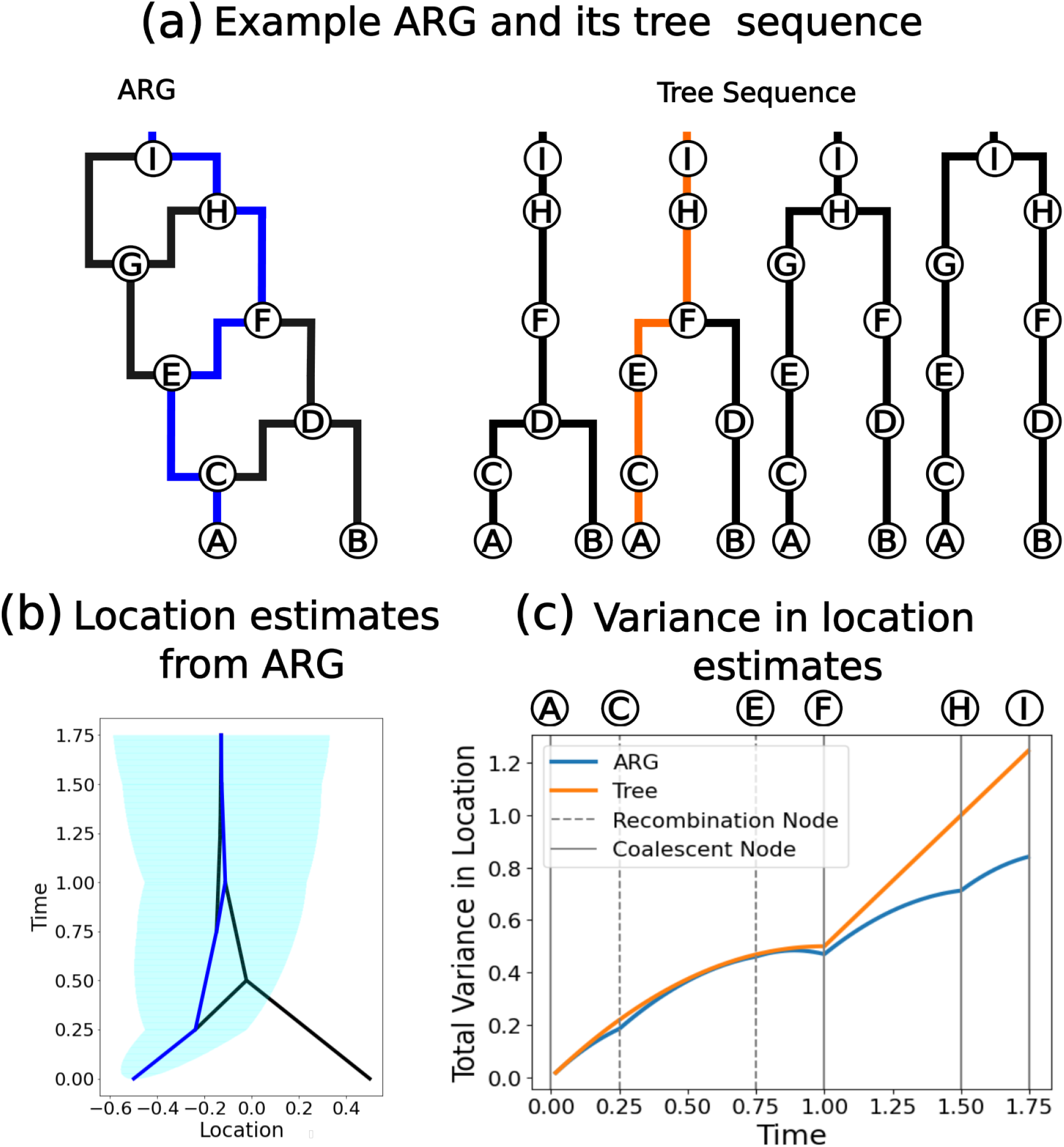
Uncertainty in ancestor locations. (a) Example ARG and the equivalent tree sequence representation with the same path highlighted. (b) The most likely location estimates (lines) and the 95% confidence interval of the blue path (shading) using the ARG. (c) The variance in location estimates, assuming *σ*^2^ = 1, along the highlighted path using the full ARG (blue) and using just the single tree (orange).

### 3.2 Dispersal rate

For a given number of samples, each additional recombination event in the ARG leads to a more clustered distribution of sample locations in the forward-in-time model. This means that, for a given set of sample locations, each additional recombination event in the ARG increases the dispersal estimate. We demonstrate the increased clustering of sample locations under the forward-in-time model in Figure 4. Focus on the variance in the location of the recombination node, *E*, and first consider the two paths to *E* (*G* → *E* and *G* → *F* → *E*) independently. The variance of the location of *E* along either path is *σ*^2^*t*, where *t*is the time between *G* and *E*, which corresponds to the variances in the location of *E* in each of the two the marginal trees. However, upon conditioning on the loop, *η*_*loops*_, the variance in the location of *E* reduces to half of its unconditioned variance, *σ*^2^*t*/2. More generally, we can treat the loop (*G* → *E* → *F* → *G*) as a Brownian bridge from *G* back to *G* and therefore calculate the variance at any point along the loop *X* as 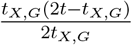 where *t*_*X,G*_ is the time between *X* and *G* (for instance, *t*_*E,G*_ = *t*) (Chow, 2009, Eq 12). Intuitively, two lineages starting at the same point are more likely to meet again if they do not wander too far away from one another. This cascades down to reduce the variance of the sample locations below recombination nodes, leading to more clustered sample locations (see Section S5 for a proof and Figure 5c for simulations).

**Figure 4:**
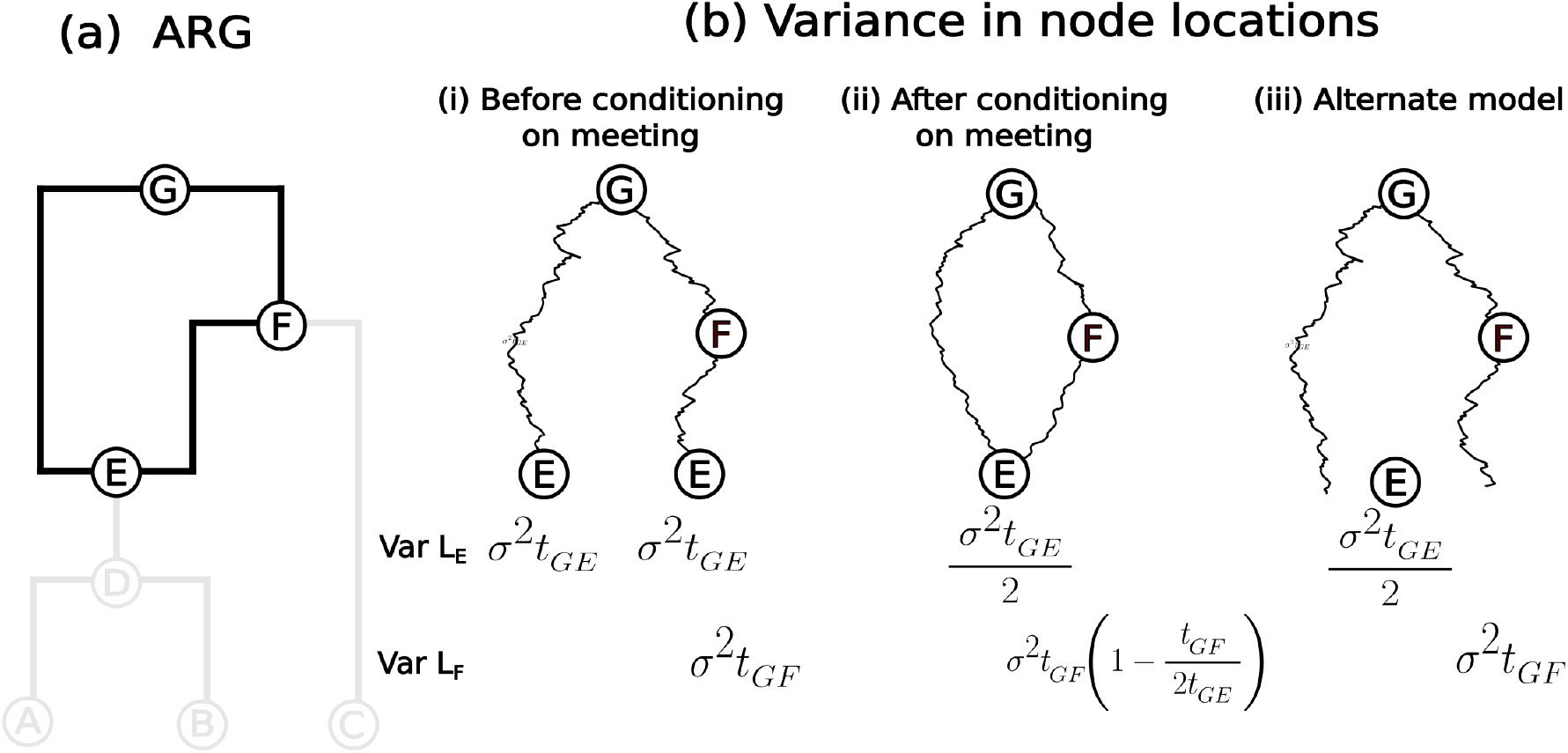
Variance in sample locations. (a) The ARG with the loop highlighted. (b) The variances of nodes E and F of the loop (i) before conditioning on the two paths along the loop meeting at E, (ii) after conditioning on them meeting and (iii) for an alternate model, where E is located at the average of the locations of its two parents.

**Figure 5:**
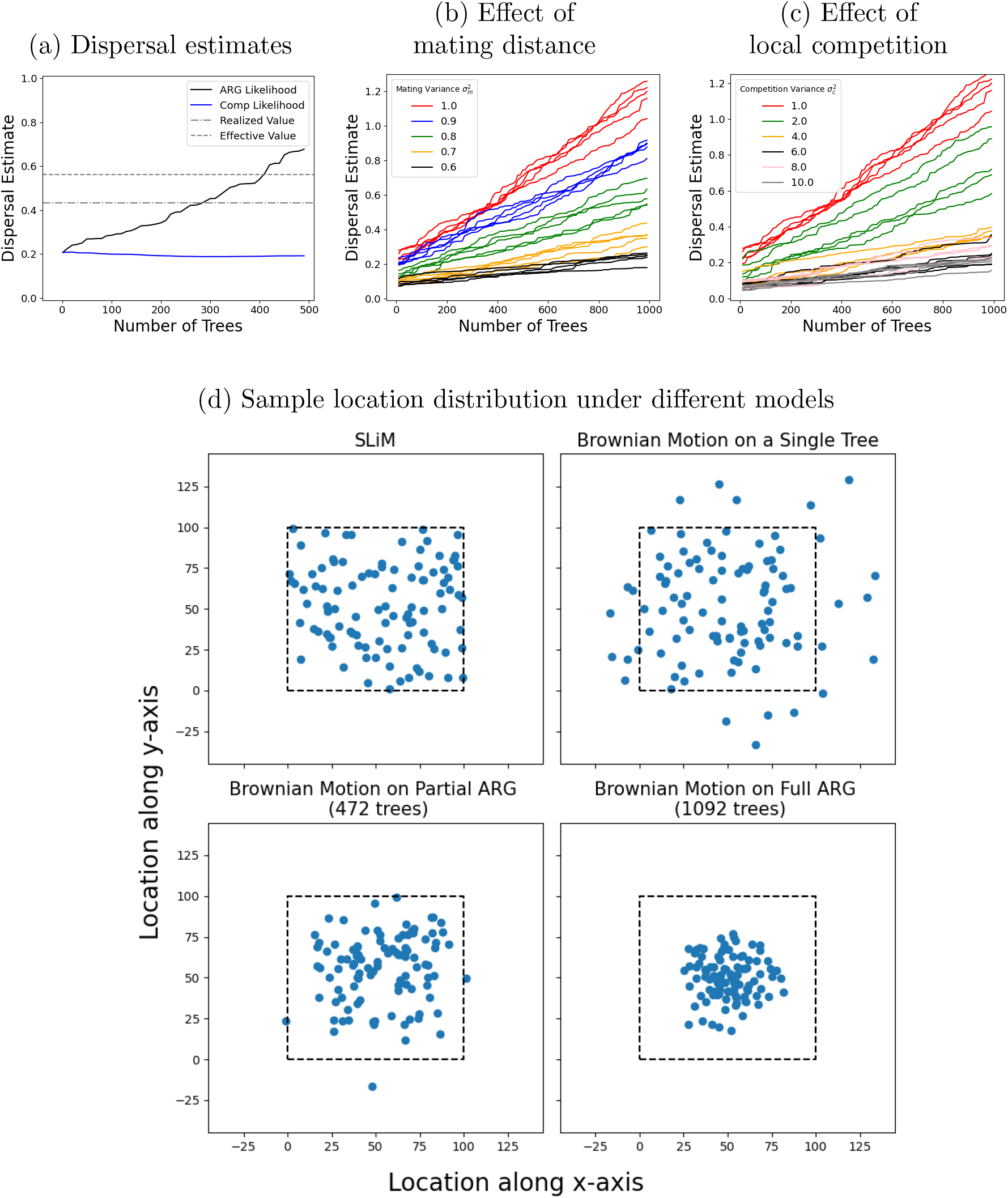
Dispersal rate accuracy. (a) Dispersal rate estimates as a function of the number of trees in the ARG used in the computation. We compare the maximum composite likelihood estimate over marginal trees (blue) and the maximum likelihood estimate from the ARG composed of the same set of trees (black). The dashed line is the effective dispersal rate (averaged over maternal and paternal variances) used in the simulation. The dashed-dotted line is the realized dispersal rate in the simulation (average squared displacement per time over all edges), which accounts for habitat boundaries. (b-c) ARG maximum likelihood dispersal estimates as a function of the number of trees colored by the variance in the (b) mating kernel and (c) competition kernel (5 replicates each).(d) The expected distribution of sample node locations under four different models. In the top left we show the sample locations from a replicate of the individual-based simulation. We then use this simulated ARG to draw sample locations from a model of Brownian motion down a single tree (top right), down a portion of the ARG (bottom left), and down the full ARG (bottom right). The dotted line is the habitat boundary of the simulations.

This clustering also occurs, though to a lesser extent, in an alternate model of Brownian motion on an ARG (Bastide *et al*., 2018), which allows the parents of *E* to be any distance apart and places *E* at their midpoint (Figure 4b-iii). Here, for a single loop, the variance of the location of the recombination node is the variance in the average of its parents locations, Var((*σ*^2^*t*_*G,E*_ +*σ*^2^*t*_*G,E*_)/2) = *σ*^2^*t*_*G,E*_/2, just as in our original model. The difference is that, unlike in the original model, the variance along the parental lineages (e.g., node *F*), and consequently the variance along lineages branching off from the loop (node *C*), are not reduced.

## 4 Testing the theory

### 4.1 Simulations

To assess the accuracy of estimates from this approach we perform individual-based two-dimensional spatial simulations using SLiM v4.0 (Haller and Messer, 2023), extending those run by Osmond and Coop (2024). Additional one-dimensional simulations and analyses are described in the supplementary materials (Section S8). Simulations include density-dependent reproduction and a finite habitat boundary.

Neither of these characteristics is captured directly under the assumptions of Brownian motion, though Osmond and Coop (2024) show that despite these differences, dispersal rates and ancestral locations can be accurately estimated under this model when applied to individual trees.

Simulations start with 10,000 individuals uniformly randomly distributed in a 100 *×* 100 unit area, with each individual being diploid for a 1 megabase chromosome. All individuals are hermaphrodites. In each generation an individual acts once as a mother, choosing its mate randomly based on distance (we assume a Gaussian mating kernel with variance 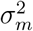) within a radius of 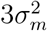. The number of offspring for each mating pair is a Poisson random variable with mean 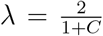, where *C* is the sum of the interaction strengths, also Gaussian with variance *σ*^2^, with neighbors within a radius of 3*σ*^2^. When there are no mates within the interaction distance no offspring are produced. Offspring are placed relative to their mother’s position with a normal random variable offset in each dimension with variance *σ*^2^. If the offset would place the offspring outside of the area, the offset is reflected off of the boundary wall and back into the area. Note that the effective dispersal rate, the expected variance in the distance between the offspring and either parent not accounting for the reflections, is given by 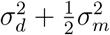 (Smith *et al*., 2023); the realized dispersal rate, accounting for the reflecting boundaries, will be lower. The locations and relationships between individuals are recorded in a `tskit` tree sequence, which is saved at the end of the simulation (Haller *et al*., 2019). Unless otherwise stated, the tree sequences are simplified to remove nodes that do not affect the broader topology of the graph and the ARG is chopped at 2000 generations in the past.

### 4.2 Dispersal rate

We first estimate dispersal rates from the simulated ARGs using Equation 6 and compare these estimates to the effective and realized dispersal rates from the simulations.

We also compute the average dispersal rates calculated over the individual trees, which is identical to the composite likelihood method of Osmond and Coop (2024) without importance sampling. Consistent with previous work (e.g., Ianni-Ravn *et al*., 2023; Kalkauskas *et al*., 2021), the dispersal estimate from the composite likelihood over trees underestimates the simulated values (blue curve in Figure 5a) due to habitat boundaries. The average dispersal estimate over all trees stabilizes as we incorporate more trees (i.e., more of the chromosome). In contrast, the dispersal estimate from our ARG likelihood systematically increases as we include more trees, starting as an underestimate but eventually leading to an overestimate of the true dispersal rate (black curve in Figure 5a).

To understand the cause of this increase in the dispersal rate estimates under our model, recall that the displacements along edges in a loop are constrained as both sides of the loop need to come back together again at a recombination node. As we saw in the previous section, for a fixed dispersal rate the forward-in-time model produces a more clustered distribution of sample locations when there are more recombination nodes (Figure 5c). Consequently, for a given set of sample locations, incorporating more trees into the ARG (and therefore more recombination nodes) must necessarily lead to a corresponding increase in the estimated dispersal rate. Further, note that as long as we model movement along the edges of the loop by independent processes and condition on them meeting at the recombination node, we will observe similar behavior in dispersal estimates. Therefore, the problem of loops is not solely associated with Brownian motion.

To confirm that conditioning on lineages meeting at recombination nodes is the core cause of the bias in dispersal rate estimates, we next run simulations that are closer to Brownian motion. Our original simulations differ from Brownian motion in two main aspects: 1) they allow some distance between mates and 2) they have local density regulation. We address these separately. First, we decrease the average mating distance by reducing the variance of the mating kernel, 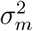 (Figure 5b). Second, we make density regulation more global by increasing the variance of the competition kernel, 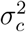 (Figure 5c). Less localized competition leads to more clumping due to the “pain in the torus” (Felsenstein, 1975) and therefore indirectly decreases the average distance between mates. As a result, both of these modifications reduce the error in dispersal rate estimates and cause them to increase more slowly with the number of trees (Figure 5b-c).

We can also investigate the cause of the dispersal rate bias by modifying our model of inference. A key assumption is that we force parent lineages to meet exactly at recombination nodes. We completely relaxed this assumption by allowing parents to mate from any distance and placing the recombination node at their midpoint (Section S4.1). However, because the variance of the recombination node is reduced to a similar extent as in the original model (Figure 4b), we still see an increase in dispersal estimates as we increase the number of trees (Section S4.1). But, unlike in the original model, the variance along the parental lineages (e.g., node *F* in Figure 4b-iii), and consequently the variance along the lineages branching off from the loop (node *C*), are not reduced. This reduces the rate at which the dispersal estimate increases with the number of trees (Section S4.1).

### 4.3 Ancestor locations

To assess the accuracy of estimated ancestor locations (Equation 7), we estimate the location of random genetic ancestors within a simulated ARG of 1000 samples and compare to the truth. We select each genetic ancestor to locate by choosing a random sample, genome position, and time (e.g., the ancestor of sample 1 at genome position 1050bp, 200 generations in the past). We estimate a location using the ARG for the full chromosome (“ARG”), a local ARG containing 100 trees on either side of the tree at the chosen genome position (“Window”), and the local tree at the genome position (“Tree”). We also compare these estimates against those using the averaging-up approach (Wohns *et al*., 2022). For the averaging-up method, we first simplified our ARG before calculating the locations (Kelleher *et al*., 2018; Wong *et al*., 2023), as is done in practice (Wohns *et al*., 2022).

Despite using more information, we observe larger absolute error with the ARG estimates than with the tree-based approach (Figure 6a). Investigating this further, we see that ancestors are often estimated too close to the average (“center”) of the sample locations (Figure 6b). Since the forward-in-time model sees excess clustering below recombination nodes (Figure 4b-ii, Figure 5d), backwards in time there is an equivalent pull towards the center (see Fig S10). This bias towards the center becomes more severe as more trees are incorporated into the ARG. Perhaps surprisingly, we see that the averaging-up method, which uses a simplified ARG and ignores edge lengths, performs just as well on average. A modified simulation with a much larger area and sampling from the center confirms this bias is not due to reflecting boundaries (Section S7).

**Figure 6:**
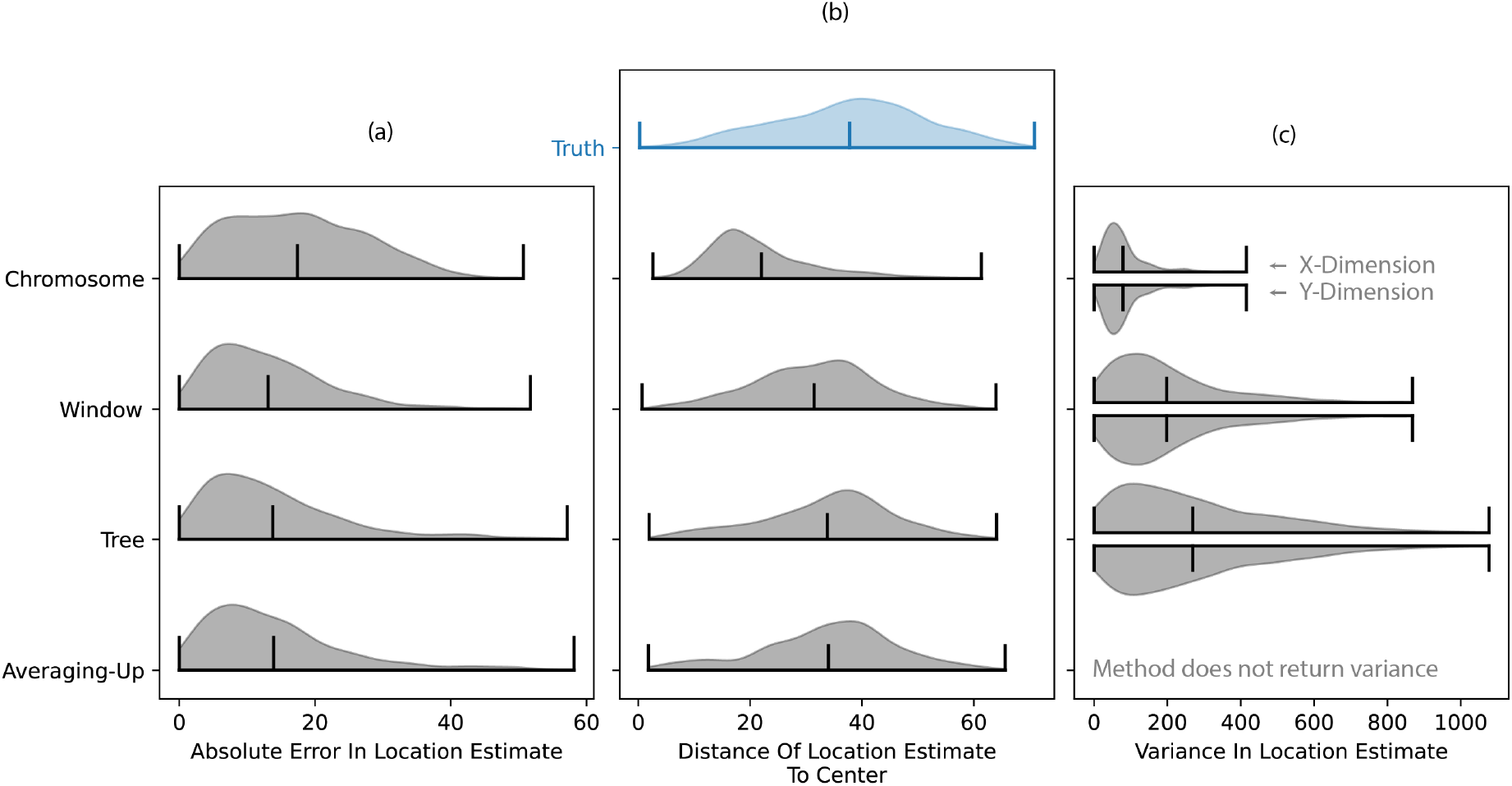
Accuracy of estimated ancestor locations. (a) Distribution of distances between the MLE ancestor locations and their true locations. Horizontal lines show the extremes and mean of each distribution. (b) Distributions of distances between the MLE ancestor locations and the average of the sample locations (“center”). The true distribution of distances of ancestor locations from the center is provided in blue for comparison. (c) Estimated variance in ancestor locations, computed using the effective dispersal rate from the simulation rather than the biased dispersal rate estimate.

Lastly, we see that the average uncertainty in location estimates gets smaller as we use more trees (Figure 6c). In practice the uncertainties would depend on the dispersal estimate but here we use the true dispersal rate to understand the unique properties of the location uncertainties, distinct from the issues of estimating dispersal. The uncertainty in location drops as we use more trees in part because each tree brings in more information. However, with more trees the model eventually becomes overconfident (Figure S9). Windowing may therefore be a promising direction forward. In these simulations at least, the number of trees in the window may be chosen so that the mean error is no worse than that from a single tree or the averaging-up method (Figure 6a) and the uncertainty is more accurate (Figure S9). However, there is currently no way of determining an appropriate number of trees to include in the window.

## 5 Promise of the approach

Acknowledging the problems illustrated above, we would like to end by illustrating the goal and promise of using an ARG-based approach for spatial reconstruction. Recombinant chromosomes show an intricate pattern of splits and merges with other sample lineages as they move through time and space. For example, in an admixed individual two regions of the chromosome will have identical spatial histories in the recent past, when they were inherited together, but very distinct spatial histories before they were brought together via recombination. To demonstrate this, we set up a simulation that starts with two geographically isolated subpopulations. Over 1000 generations, individuals in the two subpopulations disperse, intermingle, and mate in 2D. At the end of the simulation there is still a clear cline in genetic ancestry along the axis of separation between the original two subpopulations (Figure 7a). This cline is required for us to reconstruct the locations of genetic ancestors back to their original subpopulations. We then choose an arbitrary admixed individual and reconstruct the spatial histories of on either side of a recombination event. We highlight this breakpoint specifically because it separates genetic material from the two subpopulations, brought together 412 generations in the past. The ARG-based approach captures the “Y” pattern seen in the true locations, where following forward in time, the two lineages start in separate subpopulations before merging into a single path at the recombination event (Figure 7b). This pattern is not seen in estimates from individual trees as that approach does not take into account the sharing of edges between trees. Treating the two trees independently, the two lineages only come together at present day, where they meet at the sample. Other ARG-based approaches to spatial inference (Grundler *et al*., 2024; Wohns *et al*., 2022) can also capture the splitting of lineages at recombination events but do not provide measures of uncertainty, which hinders the development of future hypothesis tests.

**Figure 7:**
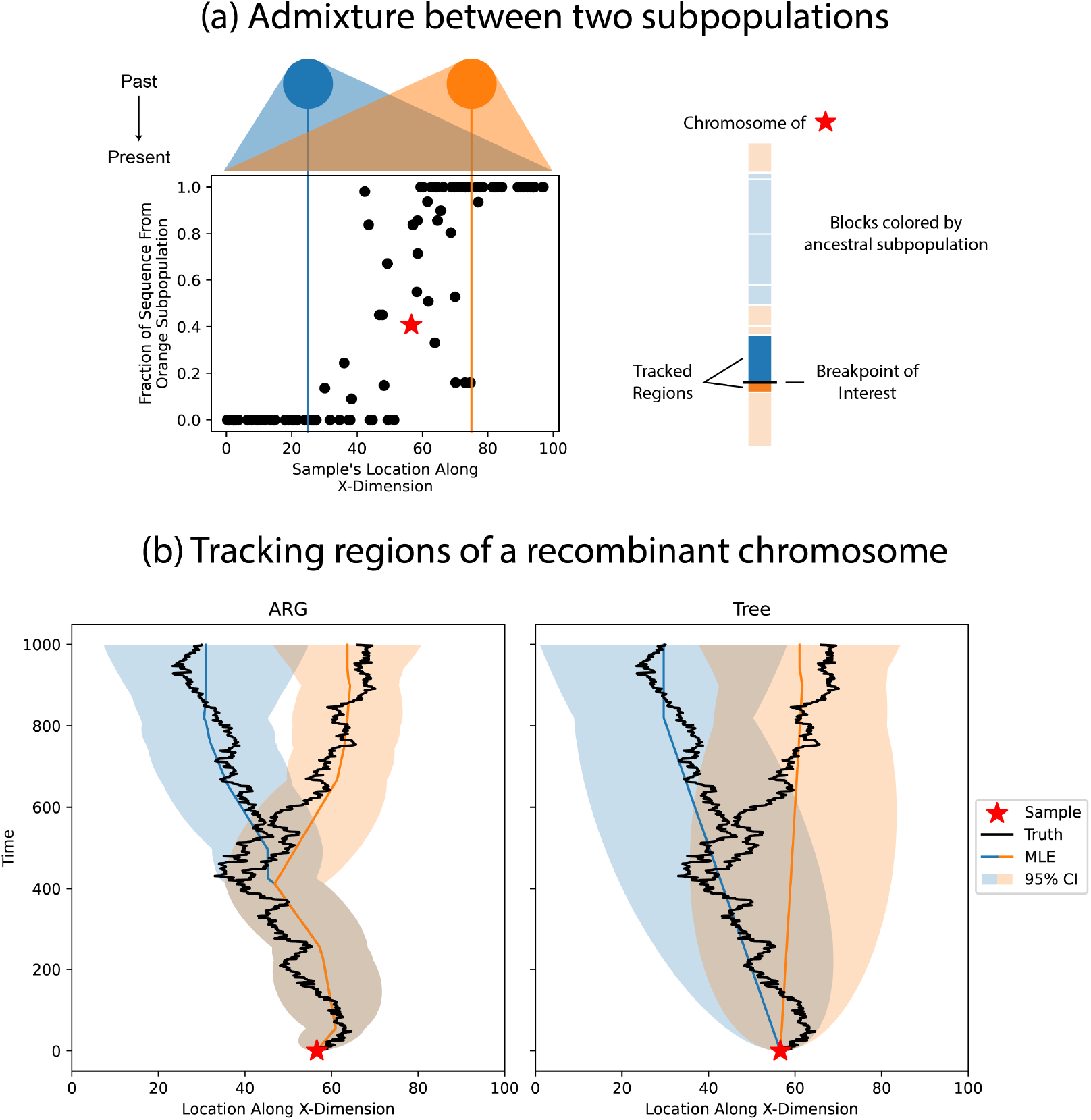
Visualizing the geographic history of admixture. We simulated two geographically isolated subpopulations dispersing into one another, leading to samples with genetic ancestors from both subpopulations. (a) At the top of subfigure is a cartoon of the dispersion of the subpopulations over time. The colored vertical lines mark the average starting position of each subpopulation along the X-dimension. As you look at samples positioned from left to right, the fraction of their sequence associated with the orange subpopulation increases. Regions of the starred sample’s chromosome are colored according to their associated ancestral subpopulation. The regions surrounding the breakpoint of interest are tracked in the following subfigure. (b) Two spatial reconstructions using the ARG for the full chromosome and the local trees immediately on either side of the breakpoint. The black lines mark the true locations of the lineages, the colored lines are the estimated most likely ancestral locations, and the shading is the 95% confidence interval around these estimates (using the effective dispersal rate from the simulation).

## 6 Data Availability

Our method is available as a Python package at https://github.com/osmond-lab/sparg. The code for all of our analyses in this paper is available in the “manuscript” branch of the GitHub repository (https://github.com/osmond-lab/sparg/tree/manuscript).

## 7 Discussion

We generalized a model of Brownian motion on trees to estimate dispersal rates and locate genetic ancestors from an ARG, and developed algorithms to do this efficiently. The full ARG provides more information than previous tree-based methods (Figure 2) and therefore estimates the location of genetic ancestors with more certainty (given the same dispersal rate; Figure 3). The great promise of such an approach is to geographically track a sample’s genetic ancestry back in time as it splits and merges via recombination and coalescence, with uncertainties. This could be particularly exciting for visualizing the divergent geography history of admixed samples (Figure 7) or locating recombination nodes of interest, e.g., viral recombination events implicated in epidemics (Ignatieva *et al*., 2022; Tamura *et al*., 2023).

Despite the promise, we discovered an interesting but unfortunate property of the model: the dispersal estimate increases monotonically as we use more marginal trees (Figure 5a). We narrowed down the root cause of this, both through simulations and mathematical proofs, to a reduction in the variance of loop node locations (and consequently that of descendant samples) under a Brownian motion model (Figure 4, Figure 5c, Section S5). Our model assumes that two lineages precisely meet at a recombination node, which is an unlikely event under Brownian motion (Etheridge, 2019), especially if they were ever far apart. This implies that the two lineages must not have dispersed very far from each other since their most recent common ancestor, pulling all nodes in and below a recombination loop closer together (Figure 5c). Consequently, given the same sample locations, each additional recombination loop increases our dispersal estimate. As each additional tree comes with an additional recombination loop, our dispersal estimate increases with the number of trees (Figure 5a). Ancestral location estimates were also shown to have a bias towards the center due to this behavior (Figure 6). Allowing the parents of a recombination node to be any distance apart reduced the rate that dispersal estimates increased with the number of trees (Section S4.1) but did not solve the problem because placing the recombination node at the average of the two parents reduced its variance.

The problem of loops poses a key challenge in utilizing ARGs for spatial inference. Despite the “pain in torus” (Felsenstein, 1975), simple models of Brownian motion often provide accurate spatial inference on trees (Bradburd and Ralph, 2019; Novembre and Slatkin, 2009). However, here we have shown that the additional and distinct problem of loops makes these simple Brownian motion models less accurate for spatial inference when extended to work on ARGs. We are therefore still without an analytically tractable (or at least computationally efficient) model that can utilize the complete information encoded in an ARG and provide accurate ancestor location estimates with confidence intervals. From the insights gained from this project, we highlight some ways forward given this problem of loops.

One way to have less biased parameter estimates is to use a more accurate model. An obvious choice is the spatial Λ-Fleming-Viot (SLFV) process (Barton *et al*., 2010a,b). By modeling local density dependence (which allows two lineages of a loop to be non-independent even before conditioning them on meeting) and non-zero mating distance (relaxing the meeting condition) this approach gives more accurate estimates than simple Brownian motion when applied to trees (Kalkauskas *et al*., 2021). The SLFV model can be extended to incorporate recombination between lineages that are not at the exact same location (Barton *et al*., 2010b; Etheridge and Véber, 2012). One of the key steps in utilizing the SLFV model for inference with ARGs is to extend the model to incorporate recombination along a continuous genome (as opposed to the two loci; Etheridge and Véber (2012)) which, while promising, is a difficult problem. Further, the complexity of this model makes it computationally intensive for inference and therefore limited to small sample sizes, even in the case of no recombination (Guindon *et al*., 2016). It may then be helpful to think of ways to develop more tractable and scalable models. In doing so, it is worth noting that the problem of loops is not specific to Brownian motion. In fact, as long as movement along edges of a loop are modeled as independent processes with restrictions coming solely due to what happens at recombination nodes, we will see an increased clustering of tip locations with the number of recombination nodes. One possible way around this is to model movement with Levy processes that are not independent to begin with, for instance compound Poisson processes, which show up as limiting cases of the SLFV (Barton *et al*., 2010a; Berestycki *et al*., 2009). Another option would be to use machine learning algorithms to correct for model misspecification. This may be especially doable here where the dispersal rate estimate increases nearly linearly with the number of marginal trees (Figure 5a), so that correction only requires knowing the slope.

Brownian models of spatial movement on coalescent trees are identical to Brownian models of trait evolution on phylogenetic trees Bradburd and Ralph (2019); Neigel and Avise (1993). Phylogenetic networks, where reticulations are due to hybridization, gene flow, or introgression, potentially provide a similar structural counterpart to ARGs. Unlike in the case of trees though, there is an important difference worth keeping in mind. With spatial movement on an ARG, two parental lineages are expected to be near one another immediately above a recombination node but when modeling trait evolution on a phylogenetic network the mean trait values of two parental populations can be arbitrarily far apart when the two populations hybridize. This is the key difference between our model and that of Bastide *et al*. (2018), which models trait evolution on phylogenetic networks as we do in our alternate model (Section S4.1), where the parents of a recombination node can be any distance apart. Although this difference (parents being physically close, hybridizing populations being arbitrarily far in trait space) is the norm, there are exceptions in both situations (e.g., long-range pollen dispersal allows parents to be far apart and genetic barriers to gene flow can restrict how different parental populations can be in trait space), which strengthens the case for more crosstalk between population genetics and phylogenetics. For example, the problem of loops is not a problem for modeling trait evolution on phylogenetic networks given the correct phylogenetic network because the model is not (necessarily) misspecified. However, our results imply that the estimated rate of evolution is extremely sensitive to the number of reticulations in the network, which is known to be difficult to estimate accurately.

In conclusion, we have developed a mathematically rigorous and tractable model that uses the complete genealogical history of a set of samples to reconstruct the spatial history of their genetic ancestors, with uncertainties. While such an approach holds great promise, for example to visualize the geographic history of admixed samples, we see that our estimates are biased. We demonstrate that the problem is reduced variation at and below recombination loops under the model. This makes the ubiquitous model of Brownian motion inaccurate for spatial inference on ARGs, leaving a gap for future tractable and computationally-efficient methods to fill.

## 8 Acknowledgements

We thank Peter Ralph, Gideon Bradburd, Mete Yuksel, Chris Carlson, and the Coop lab for helpful conversations. This work was supported by the Natural Sciences and Engineering Research Council of Canada (RGPIN–2021-03207 and DGECR-2021-00114 to MO), the National Institutes of Health (NIH R35 GM136290 awarded to GC), the National Science Foundation (NSF DISES 2307175 to GC) and the Centre for Global Change Science at University of Toronto (Graduate Student Research Award to PD). Computations were performed on the Niagara supercomputer at the SciNet HPC Consortium. SciNet is funded by: the Canada Foundation for Innovation; the Government of Ontario; Ontario Research Fund – Research Excellence; and the University of Toronto.

## Supplementary material

### S1 Likelihood of sample locations

We start by assuming that the random displacements along each edge of the ARG, *B*_*edge*_, are independent. Hence, each path, from a root to a sample, gives a unique distribution for the location of a sample, even if two or more paths end at the same sample. Given we have *n*_*p*_ unique paths, we therefore get the distributions for *n*_*p*_ sample locations, 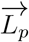. In order to get the distribution of the actual *n*_*s*_ sample locations, 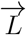, we condition on the loop conditions, *η*_loops_,

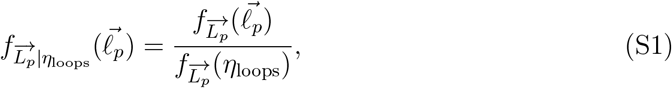

where we use the shorthand 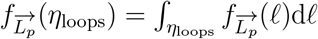 for the probability density of the loop conditions.

In Section S2 we show that the loop and path conditions are the same, *η*_loops_ = *η*_paths_. Since *η*_paths_ conditions the paths that end at the same sample to have identical locations, the numerator above becomes 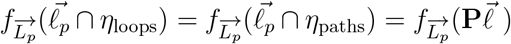 where **P** is the path-sample matrix (a *n*_*p*_ *× n*_*s*_ matrix whose *ij*^*th*^ entry is 1 if the *i*^*th*^ path ends at sample *j*). Then the distribution of path locations becomes

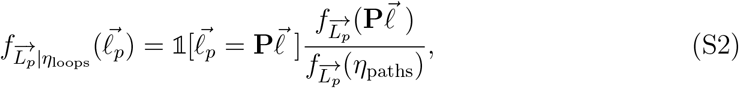

where 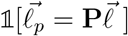 is an indicator function that is 1 if 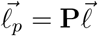 and 0 otherwise. This ensures that the probability density is 0 whenever any two paths ending at the same sample have different locations. We will now compute this distribution for the case of a single root and then do the more general multiple-root case.

#### S1.1 Single root

When the ARG has a single root, located at *μ*, the locations of path ends, 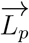, is multivariate normal with mean 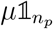 and covariance matrix *σ*^2^**S**_*p*_, where *σ*^2^ is the dispersal rate and **S**_*p*_ the path matrix (shared time between each pair of paths). Therefore the numerator of Equation S2 is

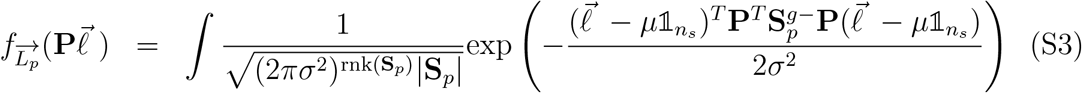

where 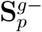 is the generalized inverse of **S**_*p*_ and rnk(**S**_*p*_) is the rank of **S**_*p*_ (which may be less than *n*_*p*_). Meanwhile the denominator is

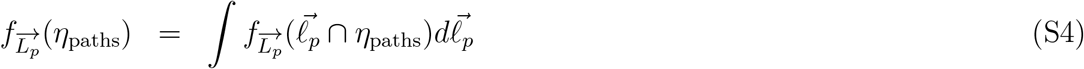

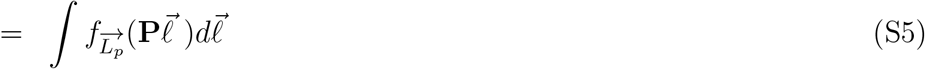

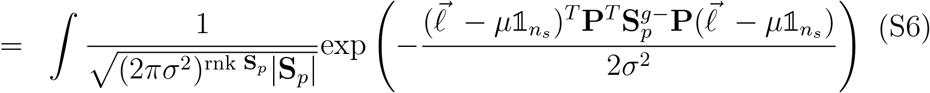

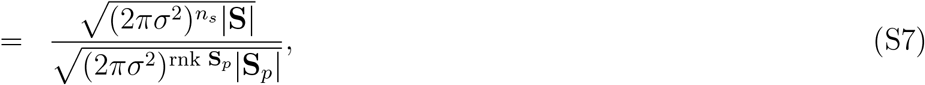

where 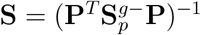 is the sample covariance matrix.

The probability density of the path locations, conditional on the loops (equivalently, paths) meeting, is then

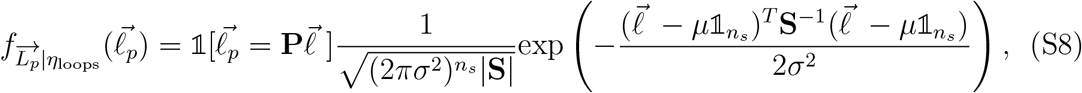

which is the probability density of a multivariate normal random variable with mean 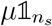 and covariance matrix *σ*^2^**S**. This is the likelihood of sample locations, 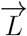, given Brownian motion down the ARG. The maximum likelihood estimates of dispersal rate and root location are then given by

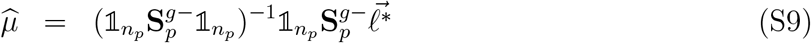

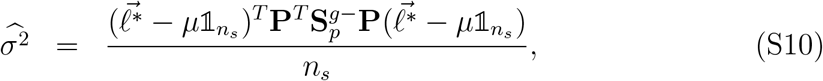

where 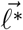 are the observed sample locations.

#### S1.2 Multiple Roots

We next want to generalize this to multiple roots, which occurs when we chop off an ARG more recently than the grand most recent common ancestor (Figure S1). Let *n*_*r*_ be the number of roots, 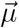 be *n*_*r*_ *×* 1 vector of root locations, and **R** be the *n*_*p*_ *× n*_*r*_ path-root matrix (the *i, j*^th^ entry is 1 if path *i* starts at root *j*, otherwise 0). Then the (unconditioned) probability distribution of the path locations, 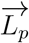, is multivariate normal with mean 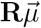 and covariance *σ*^2^**S**_*p*_. Now, as in the single root case, we want to find the distribution of the path locations conditioned on the paths meeting at the samples, *η*_paths_.

As with a single root, the numerator of wquation S2 can be written 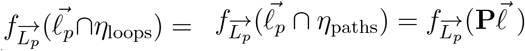, but now this is

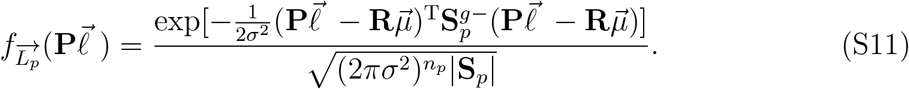

Similarly, the denominator becomes

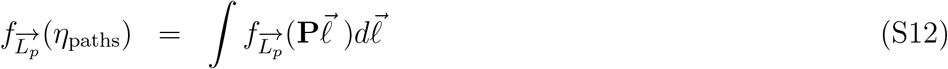

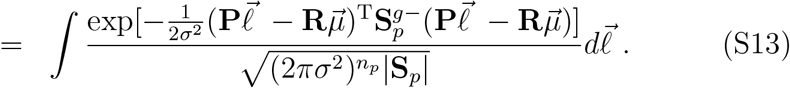

**Figure S1:**
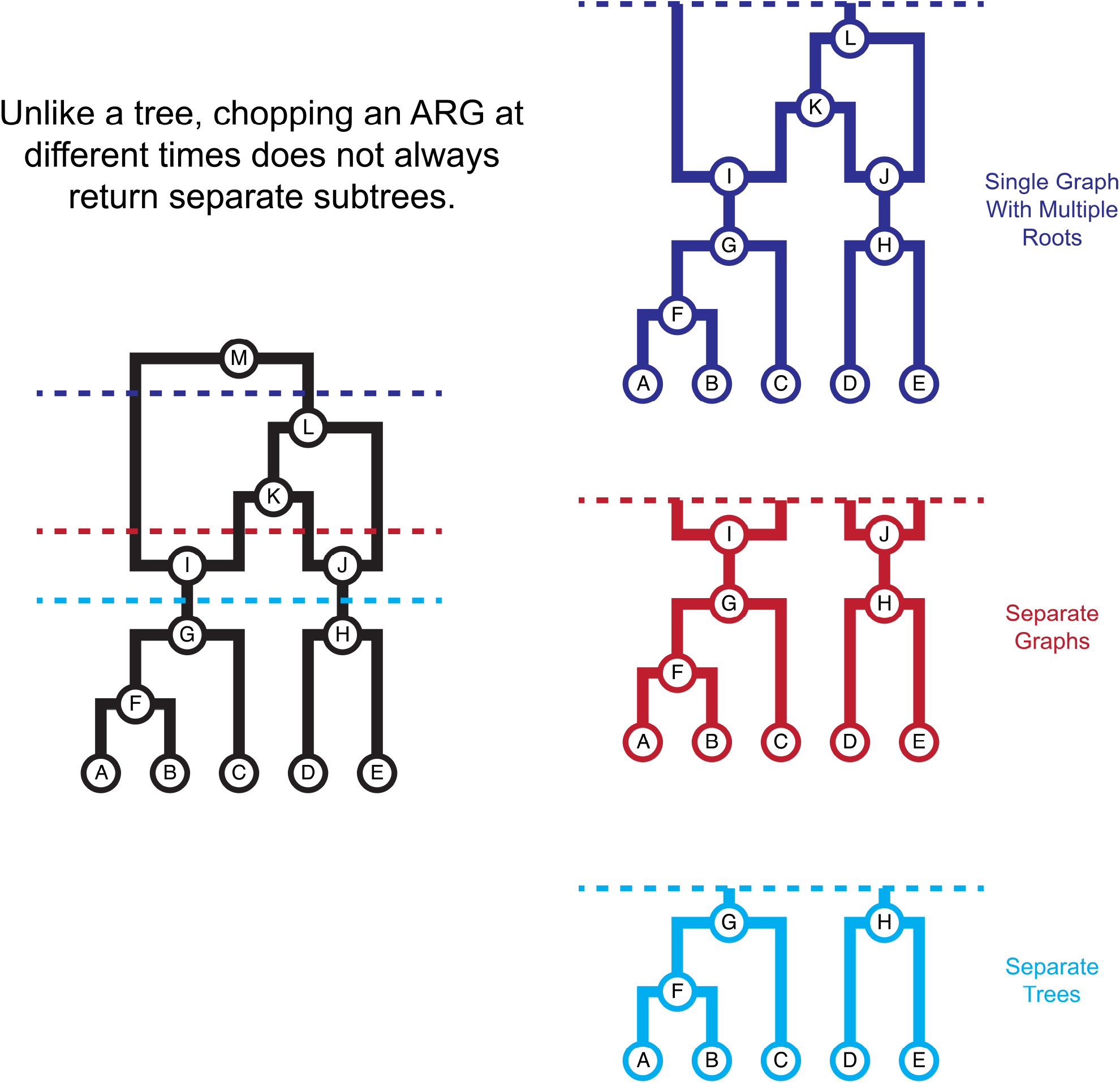
Multiple roots. Cartoon for the various scenarios that may occur when chopping an ARG below its grand most recent common ancestor.

To find this integral we multiply and divide by a constant to make the integrand a probability density for 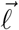. In order to do that, note that the term in the exponent can be expanded like

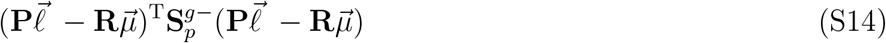

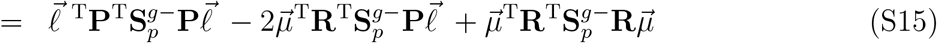

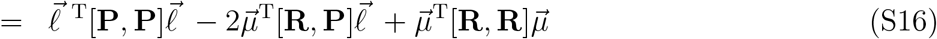

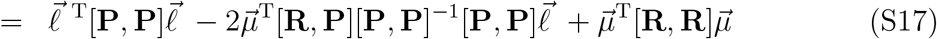

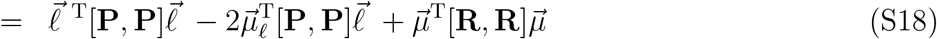

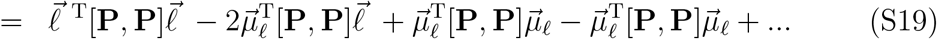

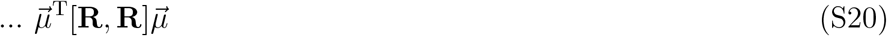

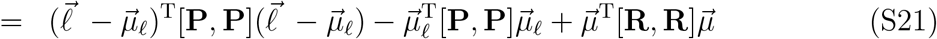

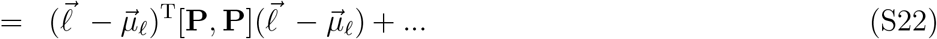

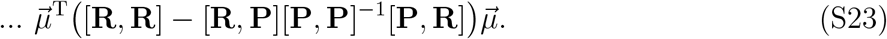

In step 1 above we have used 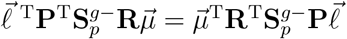, as these are 1 *×* 1 matrices and therefore are the transpose of each other. We have also used the shorthand 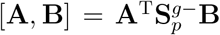 and introduced 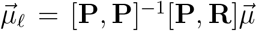, a *n ×* 1 vector which will correspond to the expectation of the sample locations (as shown below). Using this expansion, we first define

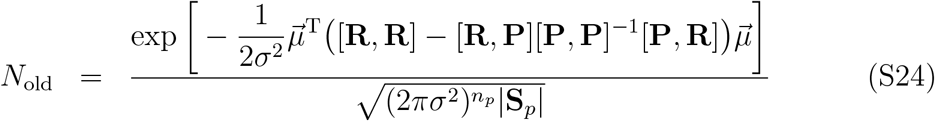

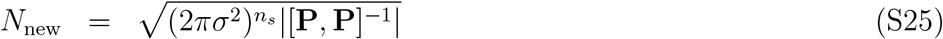

and write

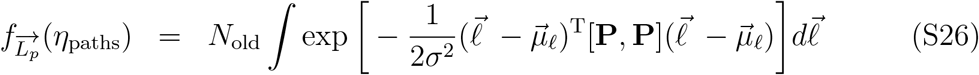

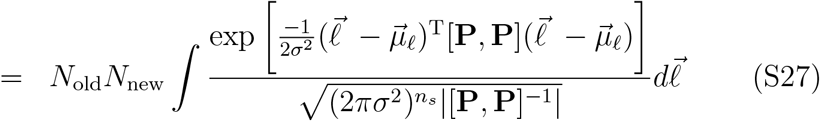

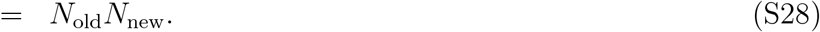

We also rewrite Equation S11 as

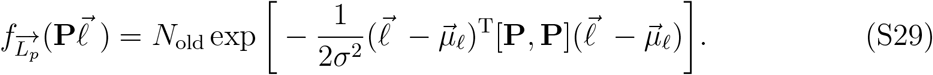

Dividing numerator by denominator, the distribution of the path locations after conditioning on the loops (equivalently, paths) meeting is

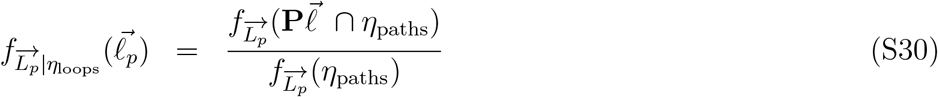

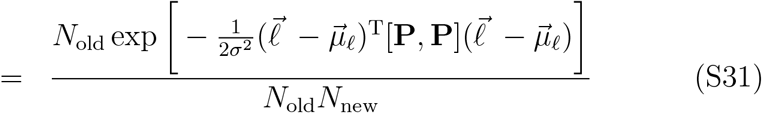

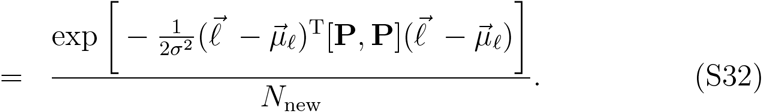

This is a multivariate normal distribution with mean 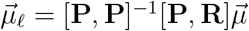 and covariance *σ*^2^**S** = *σ*^2^[**P, P**]^−1^. This is the likelihood of sample locations, 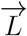, given Brownian motion down the ARG.

To derive the maximum likelihood parameter estimates, note that the log likelihood function for the parameters is given by

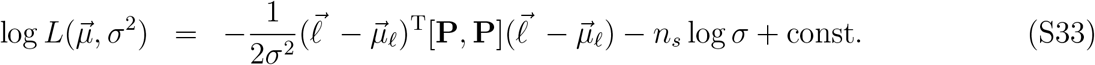

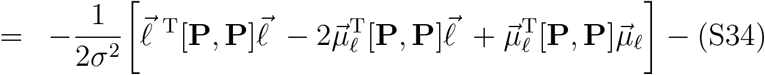

We can use this to find the maximum likelihood root locations by first differentiating the log likelihood function with respect to each root location

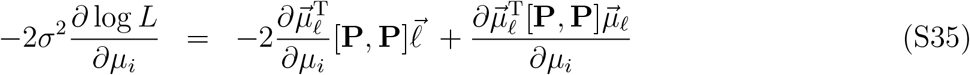

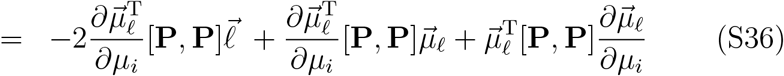

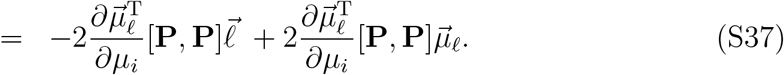

Now, since 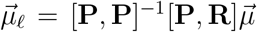 then 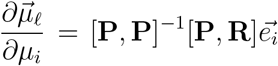, where 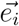 is the unit vector of length *n*_*s*_ with 1 in the *i*^*th*^ position. Therefore

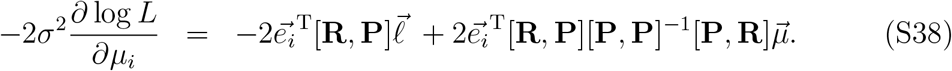

Setting the left hand side to zero for all *i* ∈ {1, 2, …, *r*} we get the maximum likelihood root locations, 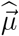, as the solutions to a system of linear equations,

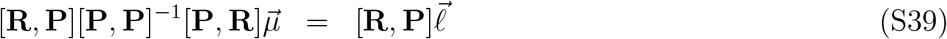

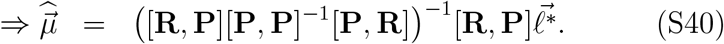

Note that using this equation to get the root locations leads to unexpected behavior. Specifically, ancestor locations rapidly move away from each other as we go back in time (Figure S2). This is probably because, forwards in time, two Brownian motions that start at different locations have the highest probability of meeting in the middle, which forces them to diverge backwards in time. To avoid this issue, we use the unconditional distribution of 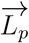 (Eq. 2, i.e., not conditioning on the paths meeting at the samples) to compute the maximum likelihood root locations,

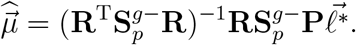

This behaves as we expected, with ancestor locations that do not diverge as strongly back in time (Figure S2), and we therefore use this method in the main text. We leave a more complete understanding of why the conditioned distribution behaves unexpectedly to future work. Note that when there is a single root the conditional and unconditional maximum likelihood root locations are equal (Figure S2) and collapse to our previously calculated MLE (Eq. S10).

**Figure S2:**
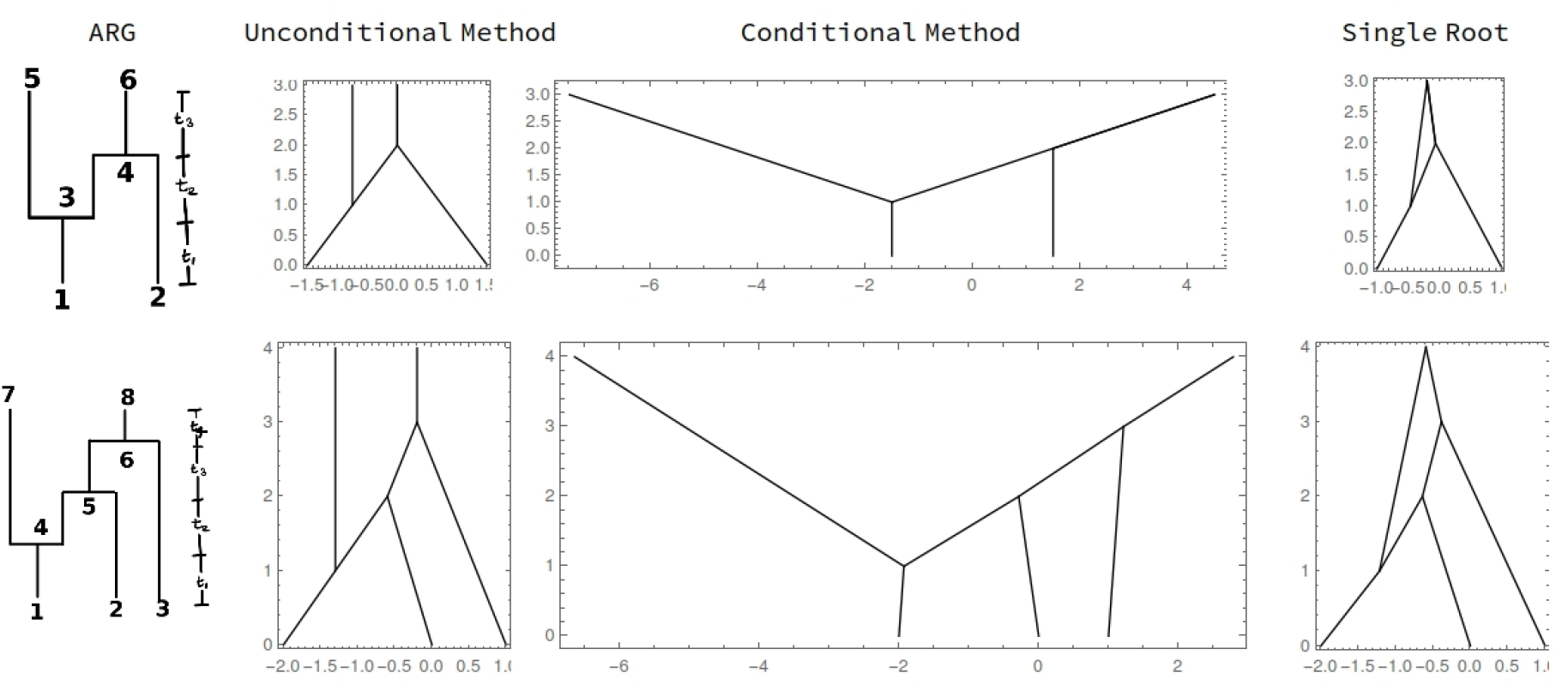
Conditional vs. unconditional ancestor locations. Inferred ancestor locations for two different ARGs (rows) with two different methods: unconditional (roots located with Equation S41) and conditional (roots located with Equation S39). When there is a single root (rightmost panel) the two methods converge.

Finally, differentiating the log likelihood (Equation S34) with respect to *σ*^2^ and setting to zero, the maximum likelihood dispersal rate when there are multiple roots is

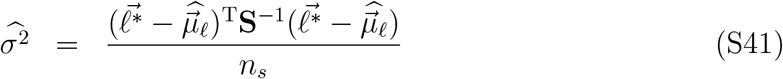

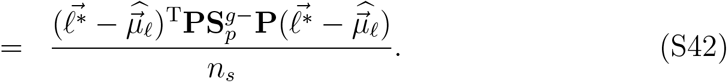

### S2 Equivalence of loop and path conditions

Here we prove that the loop and path conditions are equal, *η*_loops_ = *η*_paths_, for any ARG. Before providing a formal proof, which requires more detailed notations, we first outline the idea:

1. To prove the equivalence we need to show that for each condition in *η*_loops_ there exists an equivalent condition or set of conditions in *η*_paths_ and vice versa. In other words, for each loop we need to find a pair of paths that only differ inside that loop. And conversely, for every pair of paths from the same sample, we need to find a set of loops such that any difference in the paths belongs to one of the loops.
2. Given a loop, we find a pair of paths as follows:
3. Find the bottom (more recent) and top (more ancestral) of the loop.
4. Find a path from the bottom to one of the samples.
5. Find a path from the top of the loop to one of the roots.
6. To get the two paths, insert each of the two paths around the loop in between the two paths found above.
7. Given a pair of paths ending at the same sample, we find a set of loops as follows:
8. Start from the sample and move up one node at a time until you hit a node that is not in one of the two paths. The previous node is the bottom of the first loop.
9. Now, find the first node above the bottom of the first loop that is common to the two paths. This is the top of the first loop.
10. If the two paths are identical above the top of the first loop, then we have found the equivalent loop condition.
11. If not, repeat the steps above starting from the top of the first loop to find the next loop and so on.

#### S2.1 Formal notation

**Definition S2.1** (Directed graphs). *A directed graph, G*_*d*_, *is a two-tuple*, (*V, E*_*d*_), *where V is the finite set of vertices/nodes and E*_*d*_ ⊆ *V × V is the set of edges*.

**NOTE S2.1**. *G*_*d*_ *is a directed graph so an edge from node v to node w*, (*v, w*) ∈ *E*_*d*_, *does not necessarily imply an edge from node w to node v*, (*w, v*) ∈ *E*_*d*_. *Given an edge* (*v, w*), *we call v the parent node and w the child node. Therefore, edges are directed from a parent node to a child node*.

**Definition S2.2** (Parents). *Given a directed graph, G*_*d*_ = (*V, E*_*d*_), *with node v* ∈ *V, then ch(v) =* {*w* ∈ *V* : (*v, w*) ∈ *E*_*d*_} *is the set of child nodes of v and par(v) =* {*w* ∈ *V* : (*w, v*) ∈ *E*_*d*_} *is the set of parent nodes of v. Further*, |*par*(*v*)| *and* |*ch*(*v*)| *denote the number of parent nodes and child nodes of v*.

**Definition S2.3** (Paths). *Given a directed graph, G*_*d*_ = (*V, E*_*d*_), *with two nodes v, w* ∈ *V, a path from v to w is a sequence of vertices p* = (*v*_0_, *v*_1_, …, *v*_*n*_) *such that* (*v*_*i*_, *v*_*i*+1_) ∈ *E*_*d*_ ∀ *i* ∈ {0, 1, ‥, *n* − 1} *and v*_0_ = *v and v*_*n*_ = *w. We will say v is connected to w, denoted by v* → *w, if there exists a path from v to w. Further, we define for any* 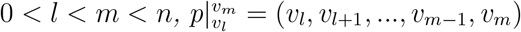,*the section of path p between v*_*l*_ *and v*_*m*_.

**Definition S2.4** (Loops). *Given a directed graph, G*_*d*_, *with two nodes v, w* ∈ *V, we say there is a loop between v and w if there exists two paths*, 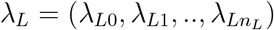 *and* 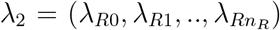, *from v to w such that* λ_*Li*_ λ_*Rj*_ ∀ *i* ∈ {1, 2, ‥, *n* − 1} *and j* ∈ {1, 2, …, *n*_2_ − 1}, *where* 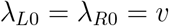 *and* 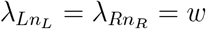.*We will denote a loop by* λ = (λ_*L*_, λ_*R*_).

**Definition S2.5** (Ancestral recombination graph). *An ancestral recombination graph (ARG) on a set of samples S is a 2-tuple*, (*G*_*d*_, *t*), *where G*_*d*_ = (*V, E*_*d*_) *is a directed graph where S* ⊊ *V, and t*: *V* → R^≥0^ *is a function associating each node with its time such that*

*1*. (*v, w*) ∈ *E*_*d*_ ⇒ *t*(*v*) < *t*(*w*)

*2. ch(s)* = ∅ ∀ *s* ∈ *S and* |*ch*(*v*)| > 0 ∀ *v* ∈/ *S*

*3*. 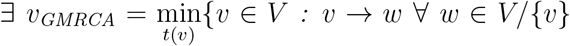 *and t*(*v*) < *t*(*w*)}. *v*_*GMRCA*_ *is called the grand most recent common ancestor (GMRCA)*.

*We say v* ∈ *V is a recombination node if* |*par*(*v*)| = 2 *and v is said to be a coalescence node if* |*ch*(*v*)| > 1.

**Definition S2.6** (SpARG). *A d-dimensional spatial ancestral recombination graph, SpARG, is an ARG and a function l* : *V* → ℝ ^*d*^ *which maps each vertex to its spatial location*.

#### S2.2 Spatial ancestral recombination graphs

We are interested in estimating the dispersal rate given a particular SpARG under a model of Brownian motion. We start by assuming that displacement along any given edge of an ARG is determined by an independent Brownian motion. We then condition on these independent Brownian motions forming the loops present in the ARG. For this we build a few more notations and definitions.

For any edge (*v, w*) ∈ *E*_*d*_, let *B*_*vw*_ ∼ *𝒩* (0, *σ*^2^*t*_*vw*_) be the random displacement along that edge under Brownian motion, where *t*_*vw*_ = *t*_*v*_ − *t*_*w*_ is the time-length of the edge. We then define the displacement function, which takes a path *p* ∈ *P* as input and returns the displacement,

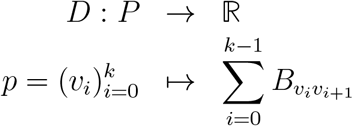

In order for these independent Brownian motions to form the loops in the ARG we need

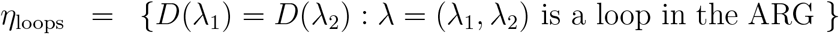

Now, let *X*_*i*_ denote the displacement of the *i*^*th*^ sample in *S, s*_*i*_, relative to the GMRCA, *v*_GMRCA_. Then the probability distribution of 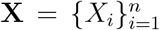, where *n* = |*S*| is the number of samples, is given by

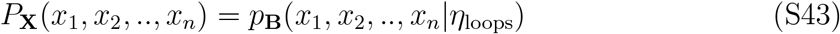

where *B*_*i*_ = *D*(*p*_*i*_) is the random variable for the displacement along a path *p*_*i*_ and 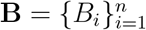.

For Equation S43 to be well defined, i.e., for it to give a single value for a given set of inputs, the value should not depend on the choice of the path from a sample to the GMRCA. To define this more formally, let 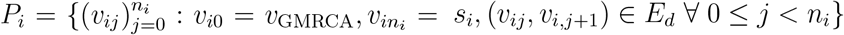 be the set of paths from the GMRCA to the sample *s*_*i*_. Now, we force the displacements along each path from the GMRCA to a given sample to be equal and call these set of conditions for all samples together as the path condition,

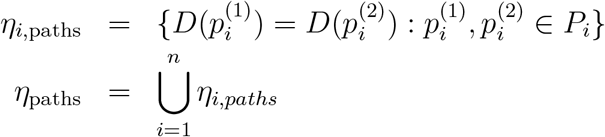

Now, as long as *η*_paths_ is true Equation S43 is well defined. We next show that *η*_loops_ = *η*_paths_, which ensures Equation S43 is always well defined.

**Lemma S2.1**. *η*_*paths*_ = *η*_*loops*_

**Proof :** [⇒] We will first show that *η*_paths_ ⊆ *η*_loops_. Therefore, we need to show that given any two paths 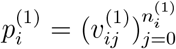 and 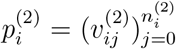 from the GMRCA to a sample *s*_*i*_, there exists loops λ^(1)^, λ^(2)^, …, λ^(*m*)^ such that

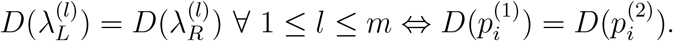

Here is how to find the loops starting from the two distinct paths. Let 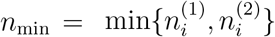 and 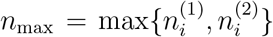. Then define 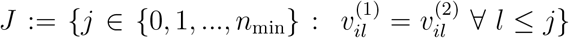 and *j*_st_ := max*J*. Therefore, *j*_st_ is the first node after which two paths start to diverge. Now, *j*_st_ < *n*_max_, otherwise we will have that 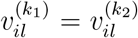 for all *l*, which would mean the two paths are identical leading to a contradiction since we started with two distinct paths. Let 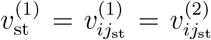. This is the start of the first loop. To find the end of this loop, let 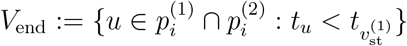. Then, the end of the loop is 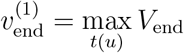. Note that 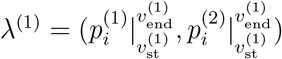 is a loop.

Now, if 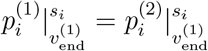, then we are done. Since everything before 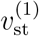 and after 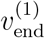 are identical in the two paths, the displacement along the two paths being equal is the same as the displacements along the two sides of the loop λ^(1)^ being equal.

If 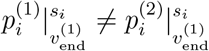, then we can repeat the above steps on 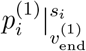 and 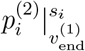, to find 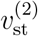 and 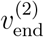 such that 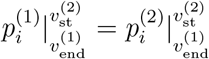 and 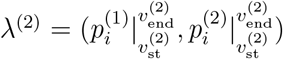 is a loop.

Keep repeating this until we have 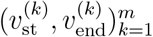 such that 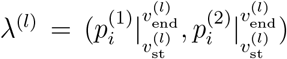 are loops and 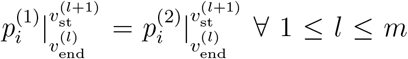 where 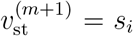. We can do this because it is a finite graph.

Therefore, the displacement along the two parts being equal is equivalent to the displacements forming the loops λ^(1)^, λ^(2)^, ‥, λ^(*m*)^. Thus, *η*_paths_ ⊆ *η*_loops_.

[⇒] Now we will show that *η*_loops_ ⊆ *η*_paths_. That is, we show that given a loop λ, there exists and a pair of paths 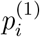 and 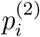 from the GMRCA to a sample *s*_*i*_ such that

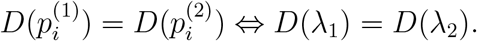

You find two distinct paths given a loop λ in the following way. Suppose λ is a loop from *v* to *w*. By definition of an ARG, *v*_GMRCA_ → *v* Let this path be 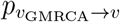. Now, if *w* ∈ *S*, then 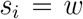 and 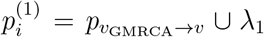 and 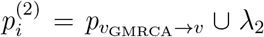 are two distinct paths from the GMRCA to the sample. Therefore, the condition for the Brownian motions to form the loop λ_1_ is the same as the displacement along 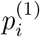 being equal to 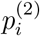.

If *w* ∈/ *S*, then we claim that there exists a sample *s*_*i*_ ∈ *S* such that *w* → *s*_*i*_. Suppose not, i.e., *w* ↛ *s*_*i*_ ∀ 1 ≤ *i* ≤ *n*. Since *w* ∈/ *S*, therefore ∃ *w*^(1)^ such that (*w, w*^(1)^) ∈ *E*_*d*_ by definition of an ARG. Now *w*^(1)^ also does not belong to *S*, otherwise we will have a vertex in *S* that is connected to *w*. Similarly, by induction we can construct {*w*^(*k*)^}_*k*∈N_ such that (*w*^(*k*)^, *w*^(*k*+1)^) ∈ *E*_*d*_ and *w*^(*k*)^ ∈/ *S* ∀ *k* ∈ N. Therefore we have infinite vertices in the *ARG* which is a contradiction. Therefore our claim has to be true. Let the path from *w* to *s*_*i*_ be 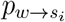, then 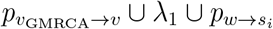 and 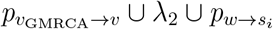 are the two required paths. Therefore, *η*_loops_ ⊆ *η*_paths_.

Therefore, we have that *η*_loops_ = *η*_paths_. Q.E.D

### S3 Minimal path matrix

The set of loop conditions, *η*_loops_, will have exactly as many conditions as the number of recombination nodes, say *k*, in the ARG. However, the size of the full paths matrix is *n*_*p*_, the total number of paths, which is greater than or equal to *k* + *n*_*s*_ (e.g., if each loop is placed alone on a terminal branch) and is bounded above by 2^*k*^*n*_*s*_ (e.g., if all loops are placed on the branch above the GMRCA). The number of conditions in *η*_paths_ is bounded by 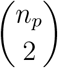, which therefore increases at least quadratically in *k* and potentially exponentially. Therefore, *η*_paths_ has multiple redundant conditions. We only need one pair of paths for each condition *η*_loops_, which only differ in one loop. We also need at least one path to each sample. Therefore, if chosen correctly, we only need *n*_*s*_ + *k* paths to calculate the correct estimates. We call the matrix of shared times of these *n*_*s*_ + *k* paths the “minimal path matrix”.

Though there are existing methods in `Python` (using the `all simple paths()` function of `networkx` package) to identify all paths from the roots to the samples, calculating the intersection between these paths does not scale well to larger ARGs, primarily due to repeated calculation of common edges across different paths. We therefore developed an algorithm, outlined below, that requires traversing each edge only once and, in doing so, have greatly sped up the calculation of **S**_*p*_.

Briefly, the algorithm entails a bottom-up traversal of the ARG starting at the sample nodes and updating the shared time matrix as we move upwards towards the roots (Figure S3). For each coalescent node visited, the algorithm calculates the edge length between that node and its parent. This is added to the corresponding cells in the shared time matrix. In addition, when we reach a recombination node (which has multiple parents), the relevant row and column are duplicated, expanding the size of the matrix and corresponding with the separation of these paths in the ARG. This keeps the size of the matrix small for as long as possible, making it more efficient.

We then add the edge length to each parent in their respective paths. Currently, the algorithm is implemented using the `tskit` package (Kelleher *et al*., 2018).

#### S3.1 Algorithm

1. Initialization
  - The shared time matrix 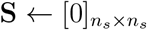, a zero square matrix of size *n*_*s*_, the number of samples. The entry of the *i*^*th*^ row and *j*^*th*^ column is denoted by *s*_*ij*_.
  - The list of paths *PL* ← [[1], [2], …, [*n*_*s*_]] with one path for each sample node.
2. Loop through every node in the ARG in time ascending order. Let *u* be the focal node. Let *I*_*u*_ be the set of indices of the paths in *PL* that currently end in *u*. Let *k*_*u*_ be the number of parent nodes. Then for each node *u*,
  a. If *k*_*u*_ = 0, *u* is the root and the loop ends.
  b. If *k*_*u*_ = 1, with parent node *v*, then do the following:
    - *s*_*ij*_ ← *s*_*ij*_ + *t*_*uv*_ for all *i, j* in *I*_*u*_, where *t*_*uv*_ is the length of edge (*v, u*). Add shared time along edge to appropriate covariance terms.
    - *PL*[*i*] ← *PL*[*i*] + [*v*] for all *i* ∈ *I*_*u*_. Extend all paths that currently end at *u* to *v*.
  c. If *k*_*u*_ = 2, with parent nodes *v*_1_ and *v*_2_, then do the following:
    - Pick one index from *I*_*u*_, say *l*.
    - *PL* ← *PL* + [*PL*[*l*]]. Duplicate the *l*^*th*^ path. Don’t update *I*_*u*_.
    - *PL*[*i*] ← *PL*[*i*] + [*v*_1_] for all *i* in *I*_*u*_. Extend all existing paths that end at *u* to *v*_1_.
    - *PL*[−1] ← *PL*[−1] + [*v*_2_]. Extend the new path formed in this step to *v*_2_.
    - 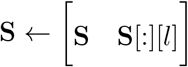. Duplicate the *l*^*th*^ column of **S**.
    - 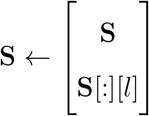. Duplicate the *l*^*th*^ row of **S**.
    - 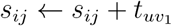 for all *i, j* in *I*_*u*_.
    - 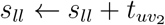

**Figure S3:**
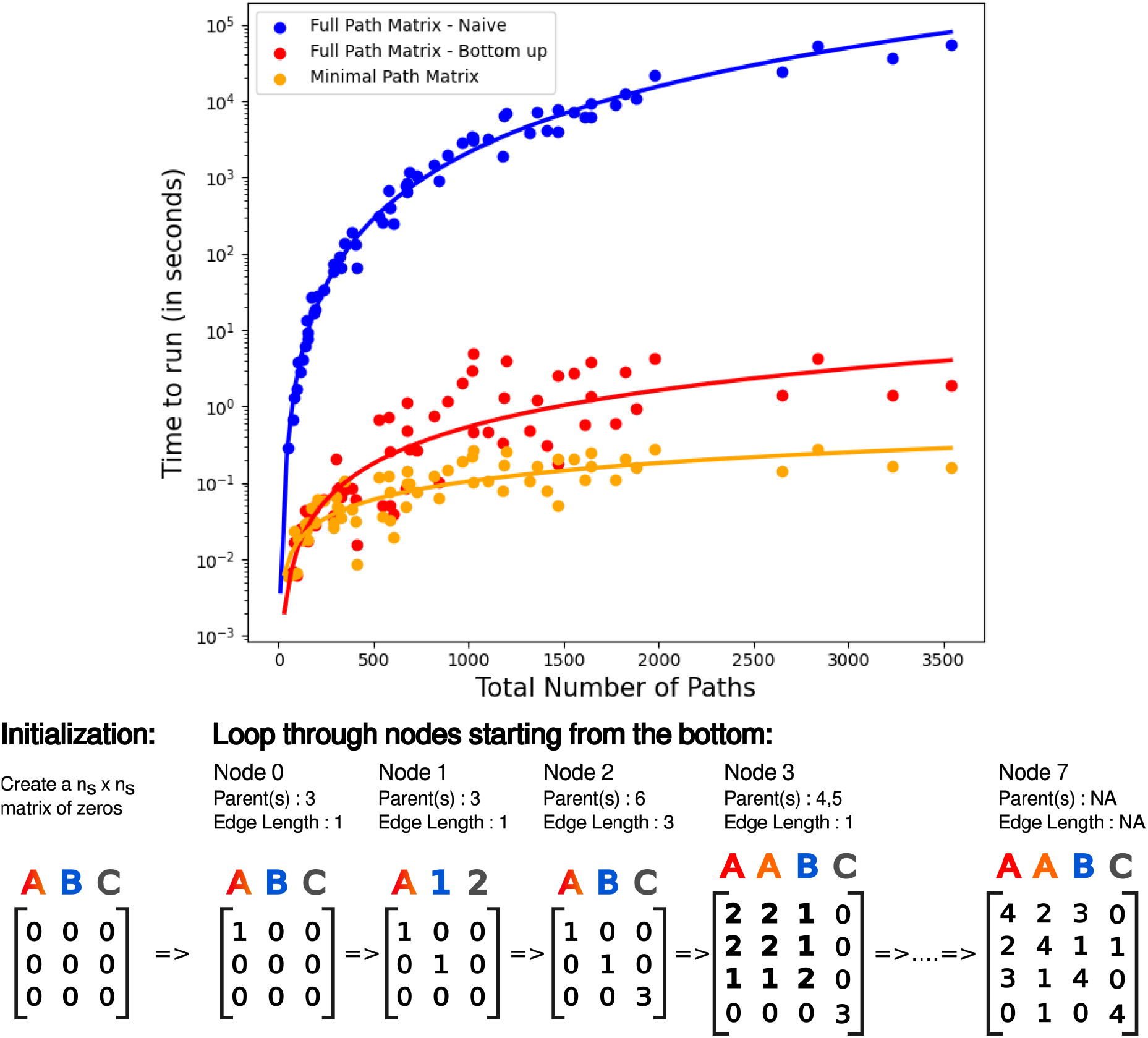
Algorithm and its benchmarks. (top) Number of seconds to compute the full path and minimal path matrices using different algorithms as a function of the total number of paths in the ARG. “Full Path Matrix - Naive” (blue) uses existing `Python` methods to compute the full path matrix. “Full Path Matrix - Bottom up” (red) instead computes the full path matrix with a single bottom-up traversal of the ARG. “Minimal Path Matrix” (orange) uses the bottom-up method to compute the path matrix for the smallest set of linearly independent paths, which is sufficient for estimating parameters of interest. Random ARGs of various sizes were generated (number of samples ranged up to 500, sequence lengths up to 5000 basepairs with recombination rate 10^−8^) using `msprime` (Baumdicker *et al*., 2022). The solid lines are the best fits under a power law. The best fit exponents for the power law are 2.946 (Full Path Matrix - Naive), 1.432 (Full Path Matrix - Bottom up) and 0.853 (Minimal Path Matrix). (bottom) Steps of our algorithm for the ARG in Figure 1.

The end result is **S**, the minimal path matrix.

### S4 Alternative models

Here we explore the dispersal estimates from two alternative models: (a) the relaxed meeting model and (b) the windowing approach. The dispersal estimates from both models are shown as a function of the number of trees used in the partial ARG in Figure S4. We briefly describe the two models and the behavior of their dispersal estimates below.

#### S4.1 Relaxed meeting model

One alternative is the relaxed meeting model, which is identical to our primary model except the parents of a recombination node need not be close to one another in geographic space (here parents means the actual parents of the recombination node, one generation back, not the parent nodes of the recombination node in the ARG). The location of the recombination node is then taken to be average of its parent’s locations. The sample matrix under this model, **S**_∞_, has been computed in Bastide *et al*. (2018) (simply set γ_*e*_ = 1/2 in their model), which we refer to for more details. Here we show how **S**_∞_ can be computed from the full path matrix, **S**_*p*_.

**Figure S4:**
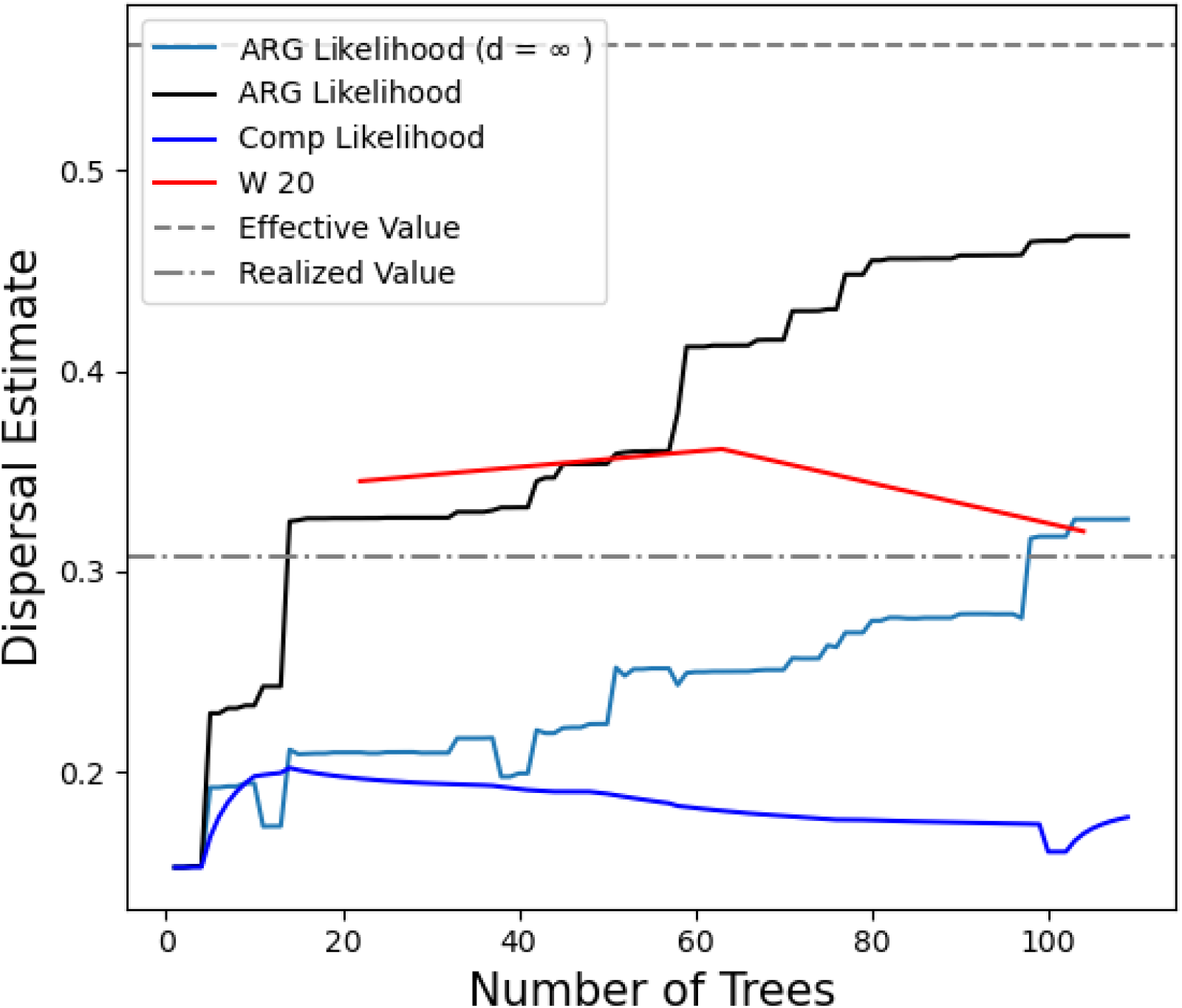
Dispersal estimates under alternative models. Dispersal rate computed from an ARG with 10 samples under different methods as a function of the number of trees. “ARG Likelihood (d =∞)” is the dispersal estimate from the full ARG under the relaxed meeting model. “W 20” is the dispersal estimate from the windowing approach which uses a partial ARG with 20 tree on each side of the focal tree. All other methods are as in Figure 5.

Let *P*_*i*_ be the set of paths from any one of the roots to sample *s*_*i*_. Then the covariance between two samples *s*_*i*_ and *s*_*j*_ is (Bastide *et al*., 2018)

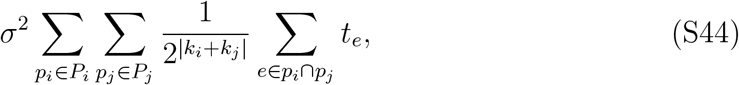

where *p*_*i*_ ∩ *p*_*j*_ is the set of common edges between the two paths and *k*_*i*_ is the number of recombination nodes along path *p*_*i*_. Let 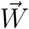 be a *n*_*p*_ *×* 1 vector which encodes the weights associated with each path. The *l*^*th*^ entry of 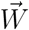 is 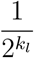. Then the sample matrix under this model is

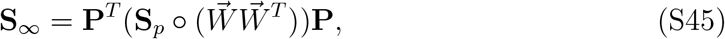

where ° is the elementwise multiplication (Hadamard product) of the two matrices. We need to extend the Bastide *et al*. (2018) to incorporate multiple roots in order to estimate dispersal rates in chopped ARGs. With multiple roots the mean of a sample location is the weighted average of the locations of all the roots it is connected to, where the weight is 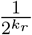 and *k*_*r*_ is the number of recombination nodes along the path from root *r*. This is given by

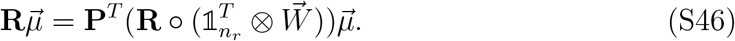

Then the maximum likelihood estimates of the root locations and the dispersal rate are

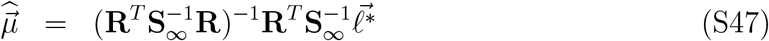

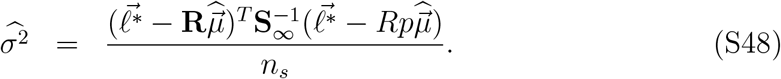

This dispersal rate estimate still increases with the number of trees (Figure S4) but the slope of increase is smaller than under our primary model and, further, we also see occasional declines. This emphasizes that the problem of loops has been reduced but not removed.

#### S4.2 Windowing approach

Another alternative is to take a windowing approach. Here we fix a window size, *w*, then build partial ARGs from disjoint groups of 2*w* trees (i.e., an ARG with trees 0 to 2*w*, an ARG with trees 2*w* +1 to 4*w*, etc.). We then take the composite likelihood of dispersal over the partial ARGs. The maximum composite likelihood dispersal estimate is then average maximum likelihood estimate over partial ARGs. For a given window size, the dispersal rate does not monotonically increase as we include more partial ARGs (Figure S4). However, larger window sizes will give larger dispersal estimates and it is not clear how to choose a good window size for a given dataset.

### S5 Recombination nodes increase clustering of sample locations under our model

Consider an ARG with *n*_*s*_ samples and *n*_*t*_ marginal trees. Now consider the partial ARG *G*_1_ over the first *k* < *n*_*t*_ trees, which has, say, *n*_*p*_ paths. Let the sample matrix for this partial ARG be **S**_1_ (of size *n*_*s*_ *× n*_*s*_) and the corresponding minimal path matrix be **S**_*p*,1_ (of size *n*_*s*_ +*k* − 1 *× n*_*s*_ +*k* − 1). Next consider the partial ARG *G*_2_ over the first *k* + 1 trees. This will have *n*_*p*_ + 1 paths. We call the corresponding sample and minimal path matrices **S**_2_ (also of size *n*_*s*_ *× n*_*s*_) and **S**_*p*,2_ (of size *n*_*s*_ + *k × n*_*s*_ + *k*), respectively. We know from Equation 3 that

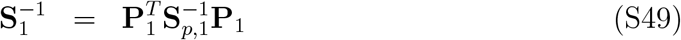

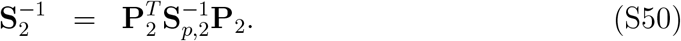

We want to show that the sample locations are more clustered for the partial ARG with more recombination nodes, *G*_2_. Let 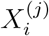, 1 ≤ *i* ≤ *n*_*s*_, *j* = 1, 2, be the sample locations distributed with covariance matrix **S**_*j*_. We are interested in the variance among these which we depict by *V* (**S**_*j*_). Since this a random variable we are interested in its expectation **E**[*V* (**S**_*j*_)],

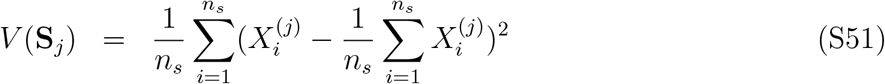

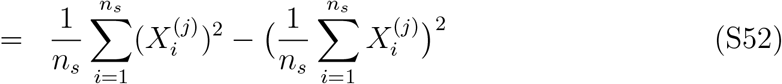

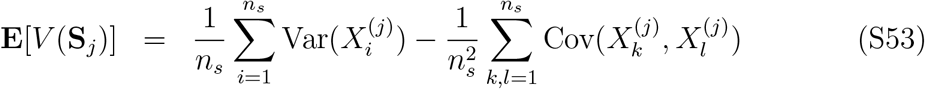

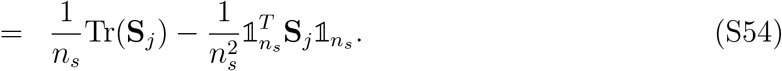

Therefore, we want to show that

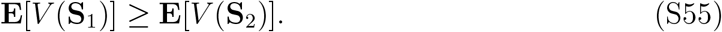

To prove this we use two properties of the variance. First, it is additive, i.e., **E**[*V* (**A** + **B**)] = **E**[*V* (**A**)] + **E**[*V* (**B**)] and second, it is positive, **E**[*V* (**A**)] > 0, for any covariance matrix **A**.

We can write **S**_*p*,2_ and **P**_2_ as an “extension” of **S**_*p*,1_ and **P**_1_, respectively. Specifically,

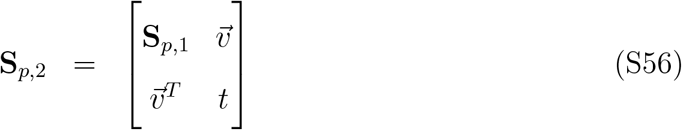

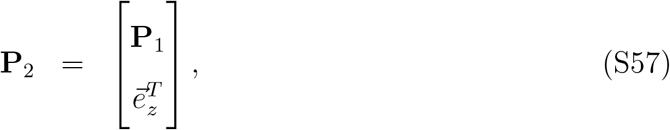

where 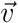 is the shared time of the new path in the minimal path set of *G*_2_ with all paths in the minimal path set of 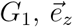 is the unit vector with all 0s except 1 at the *z*^*th*^ position, where *z* is the sample at which the new path ends, and *t*is the time from the root to the samples. Now, we can use block matrix inversion to relate 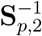 and 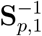,

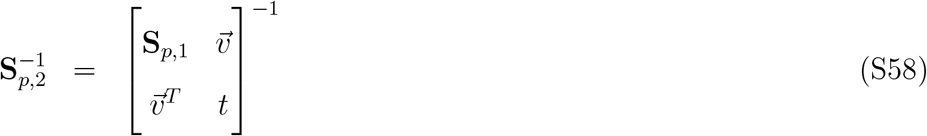

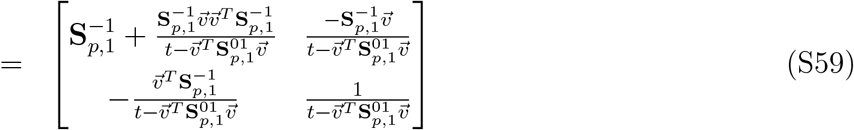

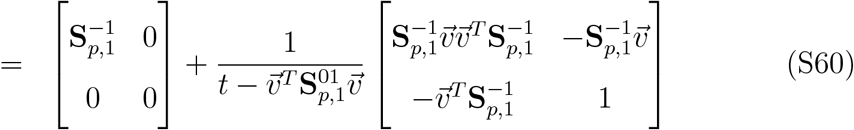

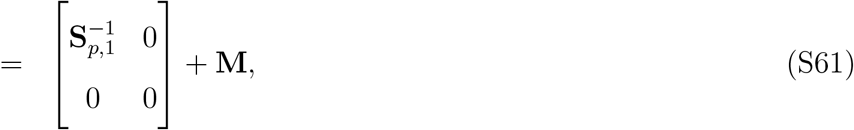

where *M* is also positive semidefinite (it can be shown that 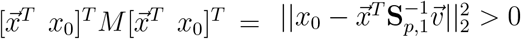 for every vector 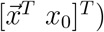. We can then relate **S**_1_ and **S**_2_,

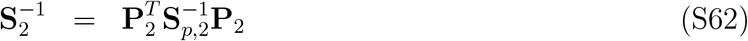

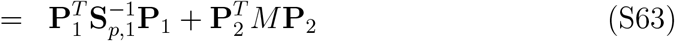

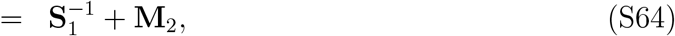

where 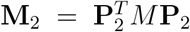 is also positive semidefinite. Now, we multiply the whole equation by **S**_1_ on the left (and right respectively) and **S**_2_ on the right (and left respectively) to get

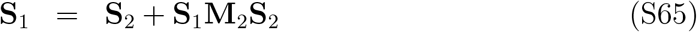

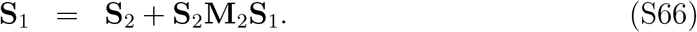

Therefore, we have that **S**_1_**M**_2_**S**_2_ = **S**_2_**M**_2_**S**_1_. Lets call this matrix **M**_3_. **M**_3_ is symmetric 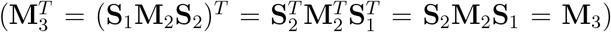 and the product of three positive semi-definite matrices. Therefore, **M**_3_ is positive semi-definite and hence a covariance matrix, which gives us

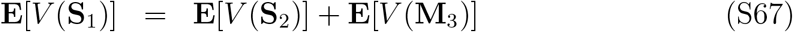

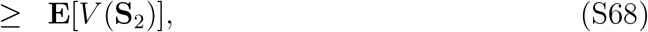

proving our statement.

We can potentially use Equation S64 to explicitly show that, for the same location of samples 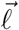, the dispersal estimate from the partial ARG with more recombination nodes, *G*_2_, is greater. To see this note that

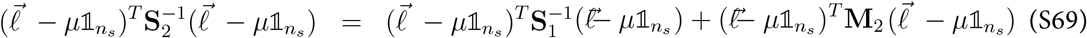

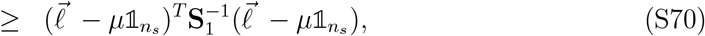

which would be a a complete proof if the root location estimate *μ* was the same for *G*_1_ and *G*_2_. Unfortunately that is not the case. Therefore, we need to prove some inequality regarding those to complete the proof, which at the moment remains elusive.

### S6 Single tree, unbounded space

To verify that our method works under our model in the absence of recombination, we simulated a single tree in unbounded space for different dispersal rates. Our estimates closely track the simulated value (Figure S5), validating our method.

Further, note that for a tree, each sample has a unique path from the root associated with it. Therefore, we have,

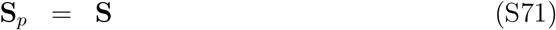

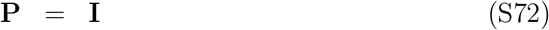

where **S** is the shared time between each pair of paths (sample lineages). Therefore, the dispersal estimate (Eq S9 and Eq S10) reduces to the well known estimates for Brownian motion on a tree,

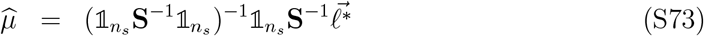

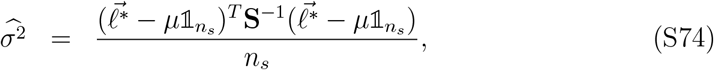

**Figure S5:**
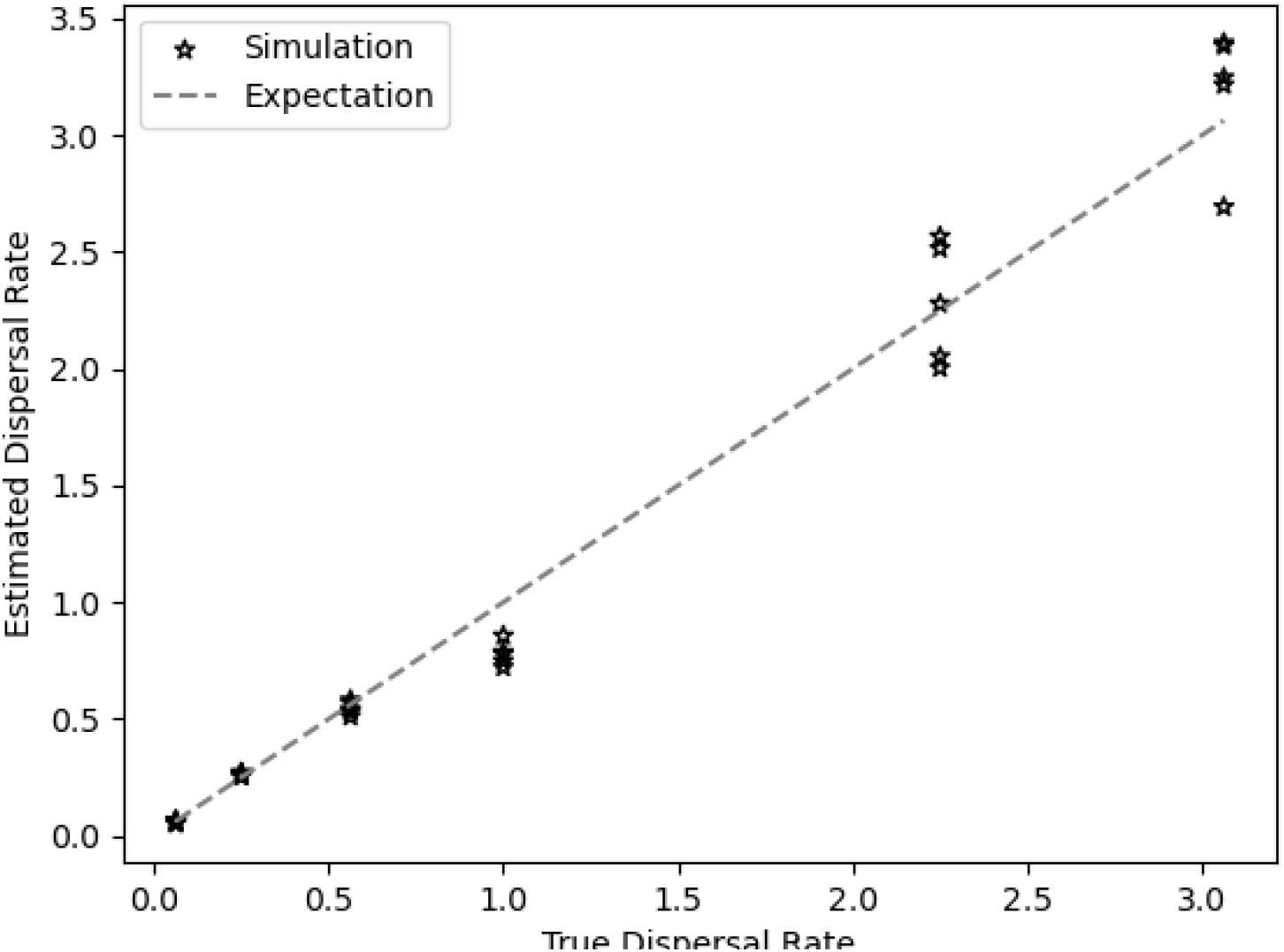
Verification of our method. Dispersal rate estimates for a tree simulated in unbounded space. Each point is a tree with 100 samples.

### S7 Boundary effects

To confirm that the reflecting boundaries in our simulations are not the main cause of the bias in location estimates, we ran the same simulations as used for Figure 6 but now in a larger area (90,000 square units versus the original 10,000 square units) but only sampled individuals from the center of the range. We expect that the shared lineages of these samples have interacted very little with the boundaries of the simulation. Even still, we observe relatively similar patterns as previously (Figure S6), in particular, a center bias. Errors were higher in this modified simulation as ancestors were able to disperse outside of the sampled range.

**Figure S6:**
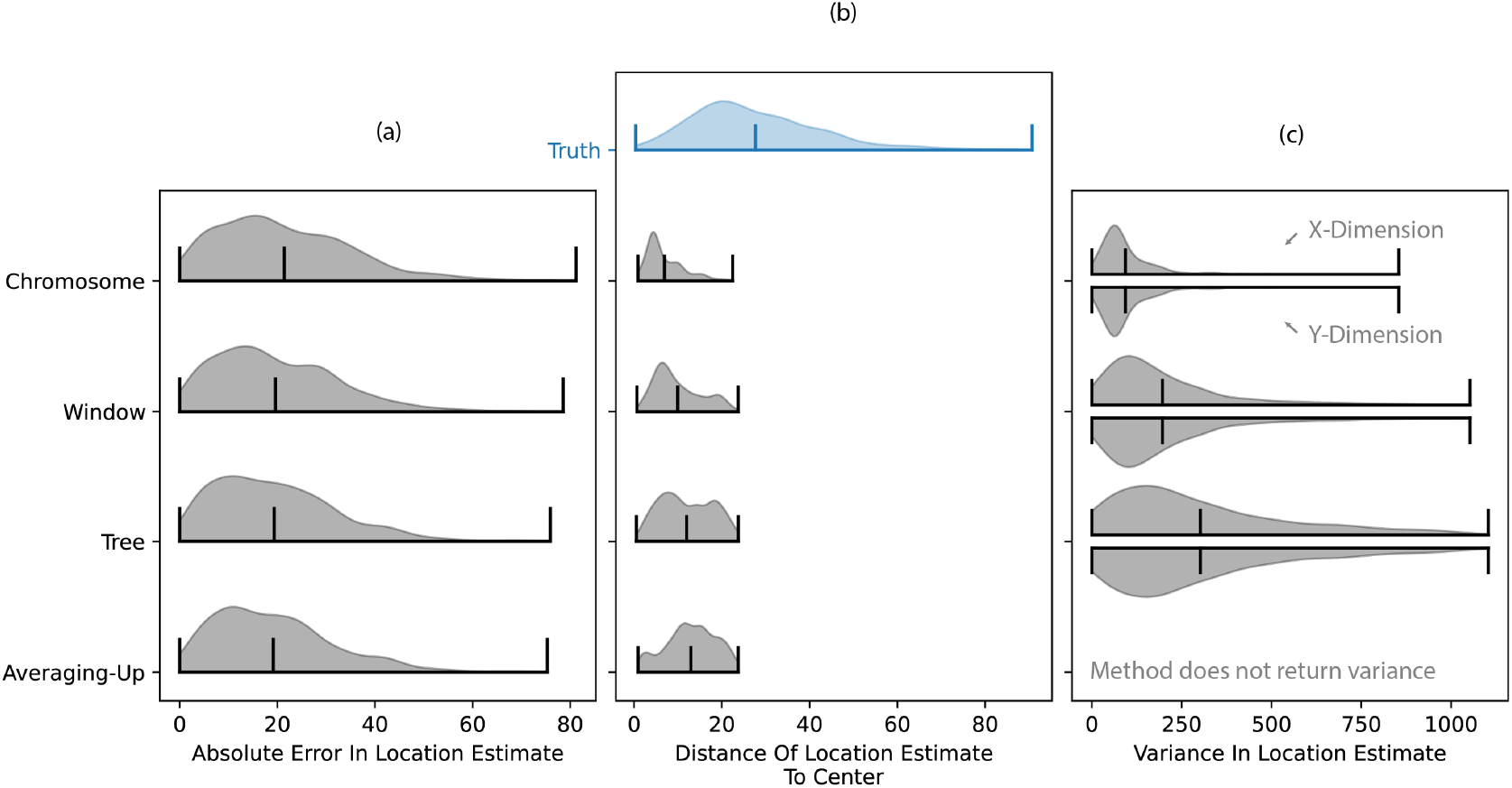
Check for boundary effects. Recreation of Figure 6 but with a modified simulation that included a larger area in which individuals could disperse. Individuals were only sampled from the center of the area; this was done to reduce any effects of the reflecting boundaries.

### S8 One-dimensional simulations

To confirm that the observed bias in our estimates is not due to a characteristic of Brownian motion in two dimensions, we reran simulations but now in one dimension. We kept the parameters consistent with the original two-dimensional simulations, but now used a simulated area that was 100 units long. Once again, we observed a higher location error when using an ARG versus the local tree, with a bias towards the center (Figure S7), and a dispersal rate that increases monotonically as more trees are included in the ARG (Figure S8).

**Figure S7:**
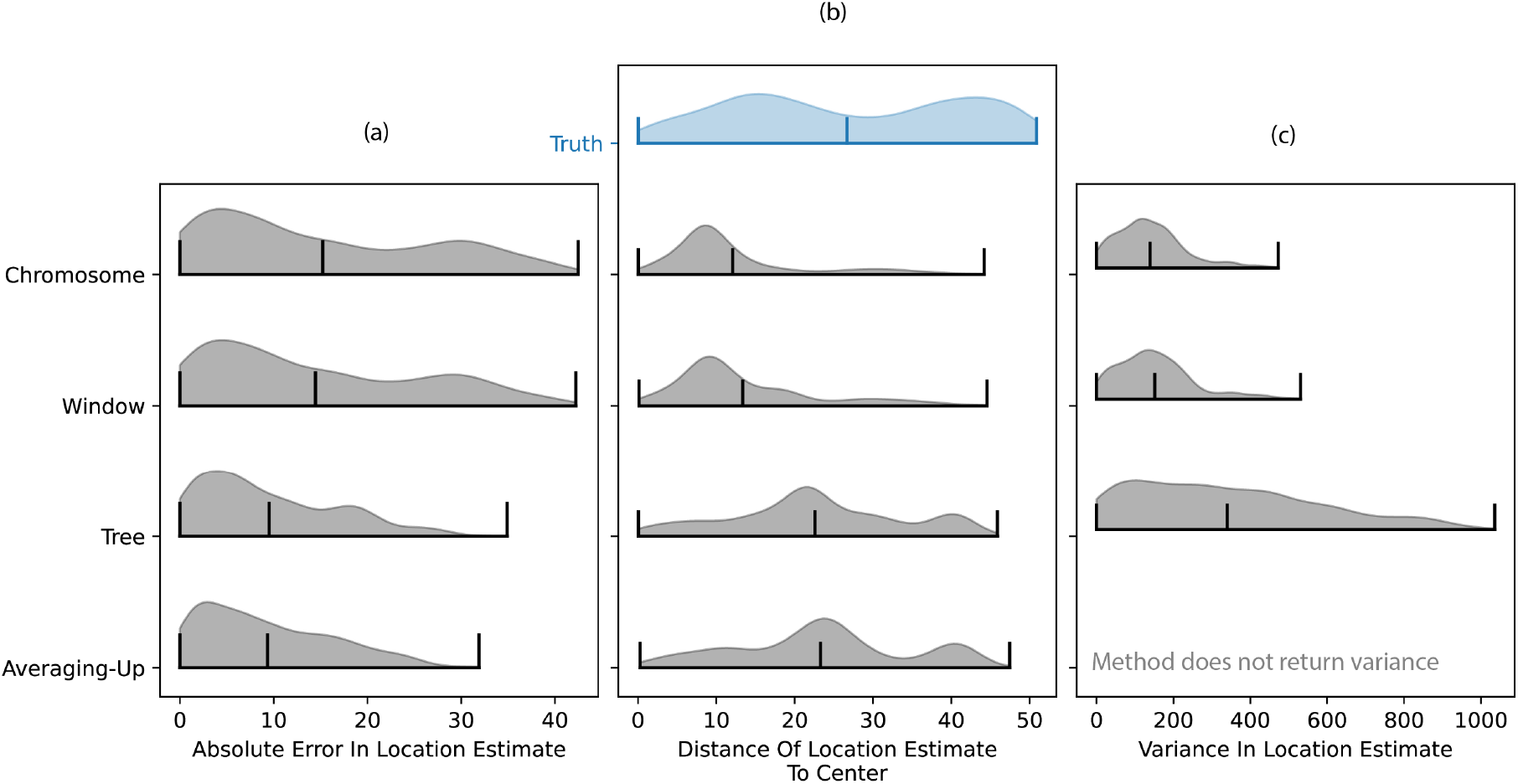
Ancestor locations from one-dimensional simulations. Recreation of Figure 6 but simulating in only one dimension. We only included 500 samples in this analysis as the population size in the one-dimensional simulation is smaller than in the two-dimensional simulation. The “Window” and “Chromosome” results are very similar here because we still used a window of 100 trees on either side of focal tree and this is relatively close to the number of trees along the chromosome.

**Figure S8:**
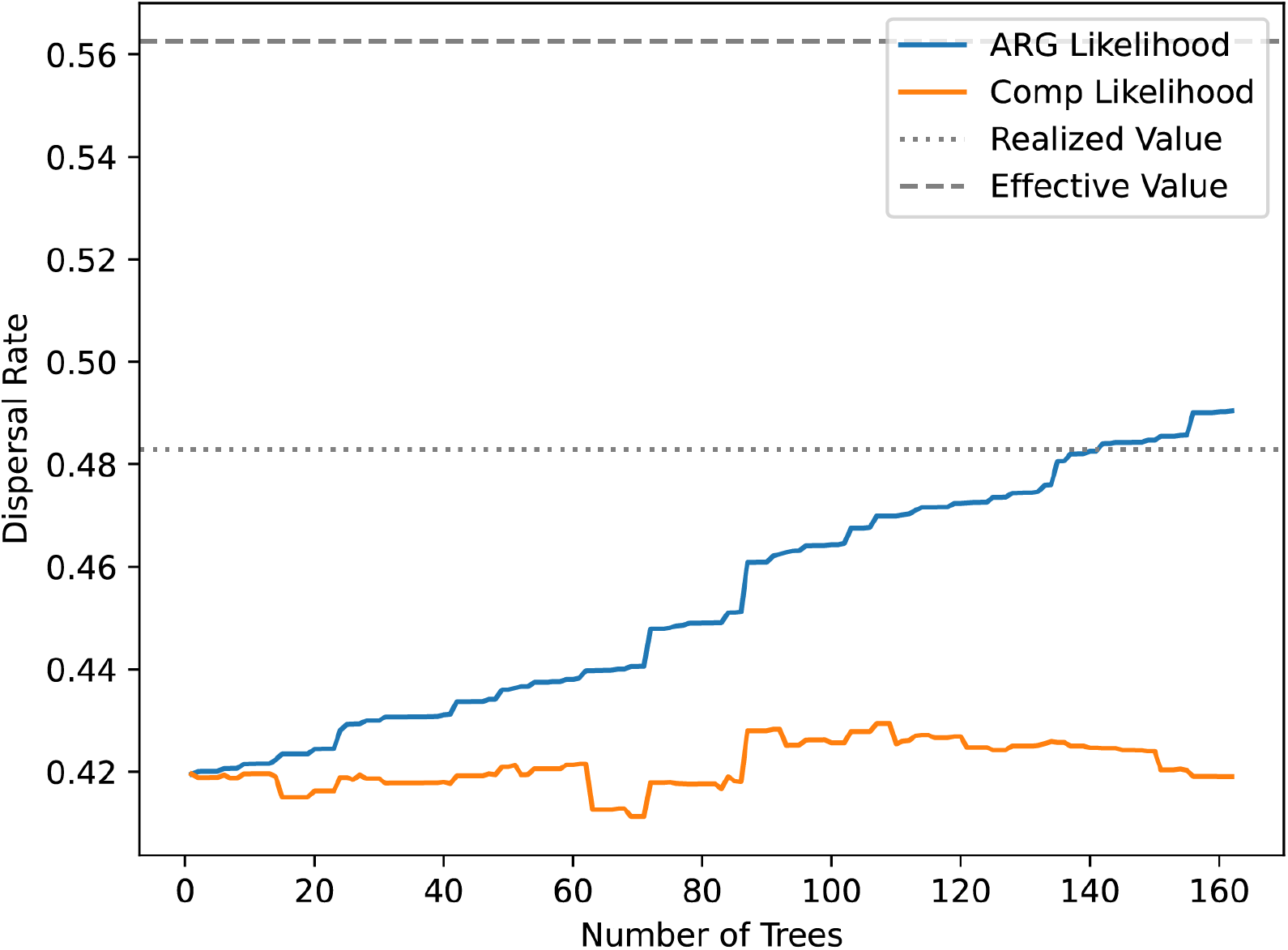
Dispersal rate from one-dimensional simulations. Recreation of Figure 5 but simulating in only one dimension (see Figure S7 for more details).

### S9 Assessing estimated uncertainty in ancestor locations

Our ARG method can be overconfident in its estimates of ancestral locations. Using the simulations from Figure 6 and the true effective dispersal rate, we created a coverage plot (Figure S9), which shows what percentage of ancestors fall within the estimates’ confidence intervals as the size of those intervals is increased. We did this using the full ARG (“Chromosome”), a window of 100 trees on either side of a focal tree (“Window”), and the focal tree (“Tree”), as in Figure 6. When using the local tree or a small window, the confidence intervals are too large. In contrast, when we use the full chromosome our confidence intervals are too small.

### S10 Ancestor location error over time

Figure S10 shows the bias in location estimates (true - estimated) color-stamped by the time. We can see that true locations greater than 50 (center of habitat) have positive error while locations less than 50 have a negative error. This is the center bias described in the main text. This center-bias gets more severe as we go back in time and as we include more trees in the ARG.

**Figure S9:**
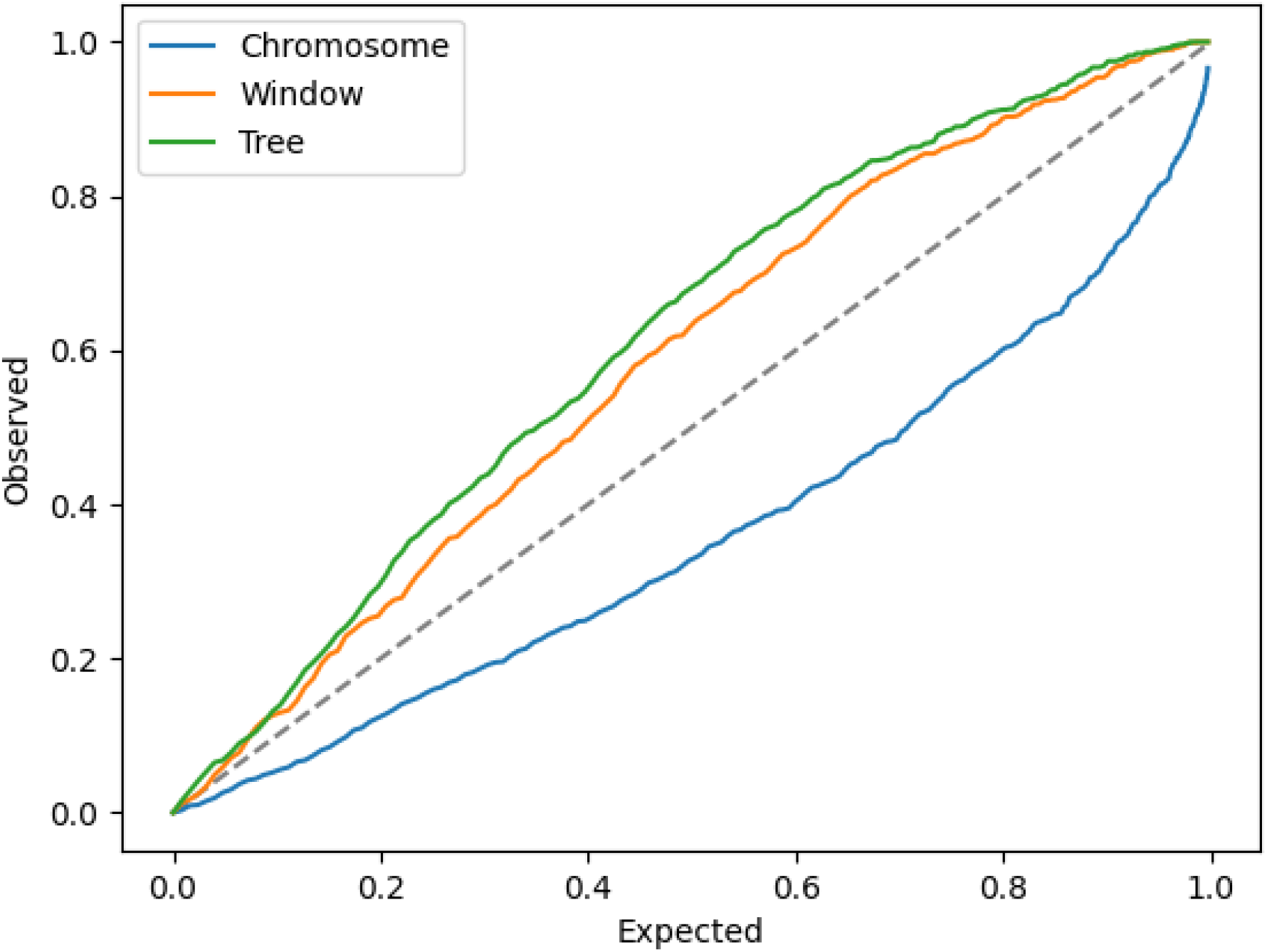
Coverage of ancestor location confidence intervals. We used the true effective dispersal rate to calculate the variance around each ancestral location estimate. We plot the observed fraction of ancestors that fall within an estimated confidence interval.

**Figure S10:**
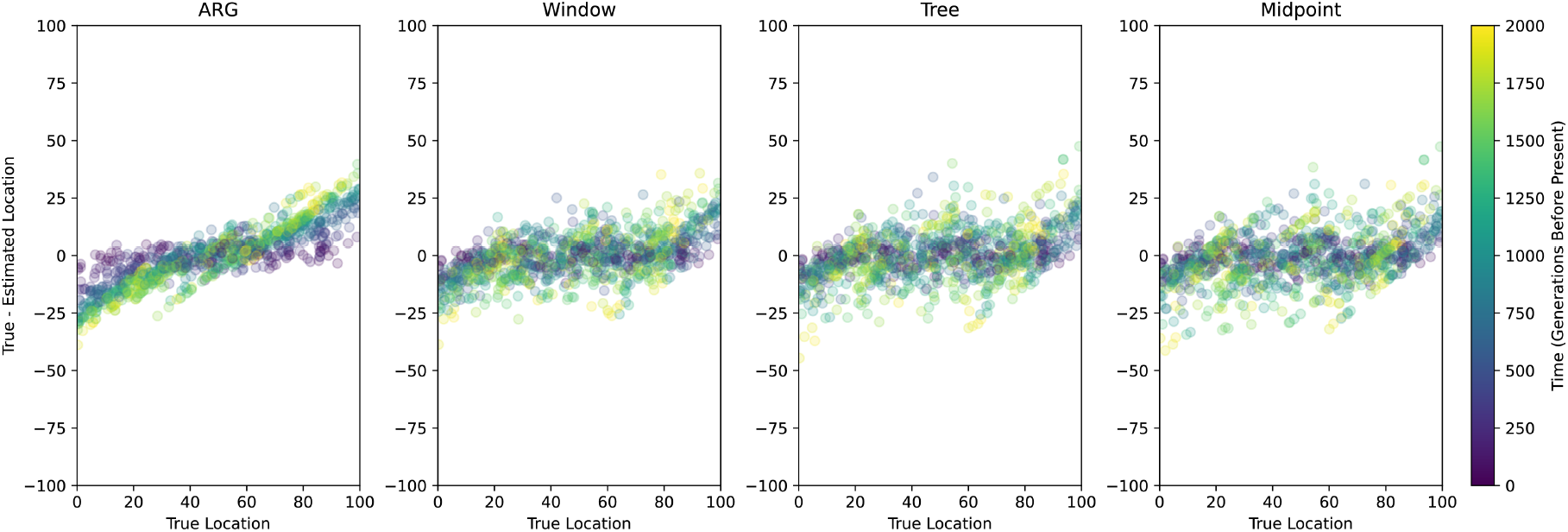
Ancestor location error by time. Error in location estimates (true - estimated locations) against the true value. The color represents the time measured backwards from present. A window of 100 trees on either side of the local tree was used for the Window panel.

